# T cell correction pipeline for Inborn Errors of Immunity

**DOI:** 10.1101/2024.09.03.610811

**Authors:** Katariina Mamia, Solrun Kolbeinsdottir, Zhuokun Li, Kornel Labun, Anna Komisarczuk, Salla Keskitalo, Ganna Reint, Frida Høsøien Haugen, Britt Olaug Lindestad, Thea Johanne Gjerdingen, Antti Tuhkala, Carolina Wieczorek Ervik, Pavel Kopcil, Nail Fatkhutdinov, Monika Szymanska, Eero Tölö, Virpi Glumoff, Janna Saarela, Trond Melbye Michelsen, Camilla Schalin-Jäntti, Johanna Olweus, Eira Leinonen, Markku Varjosalo, Eivind Valen, Timo Hautala, Martin Enge, Timi Martelius, Shiva Dahal-Koirala, Emma Haapaniemi

## Abstract

CRISPR/Cas9 gene editing technology is a promising tool for correcting pathogenic variants for autologous cell therapies for Inborn Errors of Immunity (IEI). The present IEI correction strategies mainly focus on the knock-in of therapeutic cDNAs, or knockout of the disease-causing gene when feasible. These strategies address many single-gene defects but may disrupt gene expression and require significant optimization for each newly discovered IEI-causing gene, highlighting the need for complementary platforms that can precisely correct diverse pathogenic variants. Here, we present a safe and efficient T cell single nucleotide variant (SNV) correction pipeline based on homology-directed repair (HDR), suitable for diverse monogenic mutations. By using founder mutations of Deficiency of ADA2 (DADA2), Autoimmune polyendocrinopathy-candidiasis-ectodermal dystrophy (APECED) and Cartilage Hair Hypoplasia (CHH) as IEI models, we show that our pipeline can achieve up to 80% bi-allelic editing, with resultant functional correction of the disease phenotype in patient T cells. We do not find detectable pre-malignant off-target effects or karyotypic, transcriptomic or proteomic aberrations upon profiling patient T cells with GUIDE-seq, single cell RNA sequencing, PacBio based long-read whole genome sequencing, and high-throughput proteomics. This study demonstrates that HDR-based SNV editing is a safe and effective option for IEI T cell correction and that it could be developed to an autologous T cell therapy, as the presented protocol is scalable for a GMP-compatible workflow. This study is a step towards the development of gene correction platform that targets a broad number of monogenic mutations.

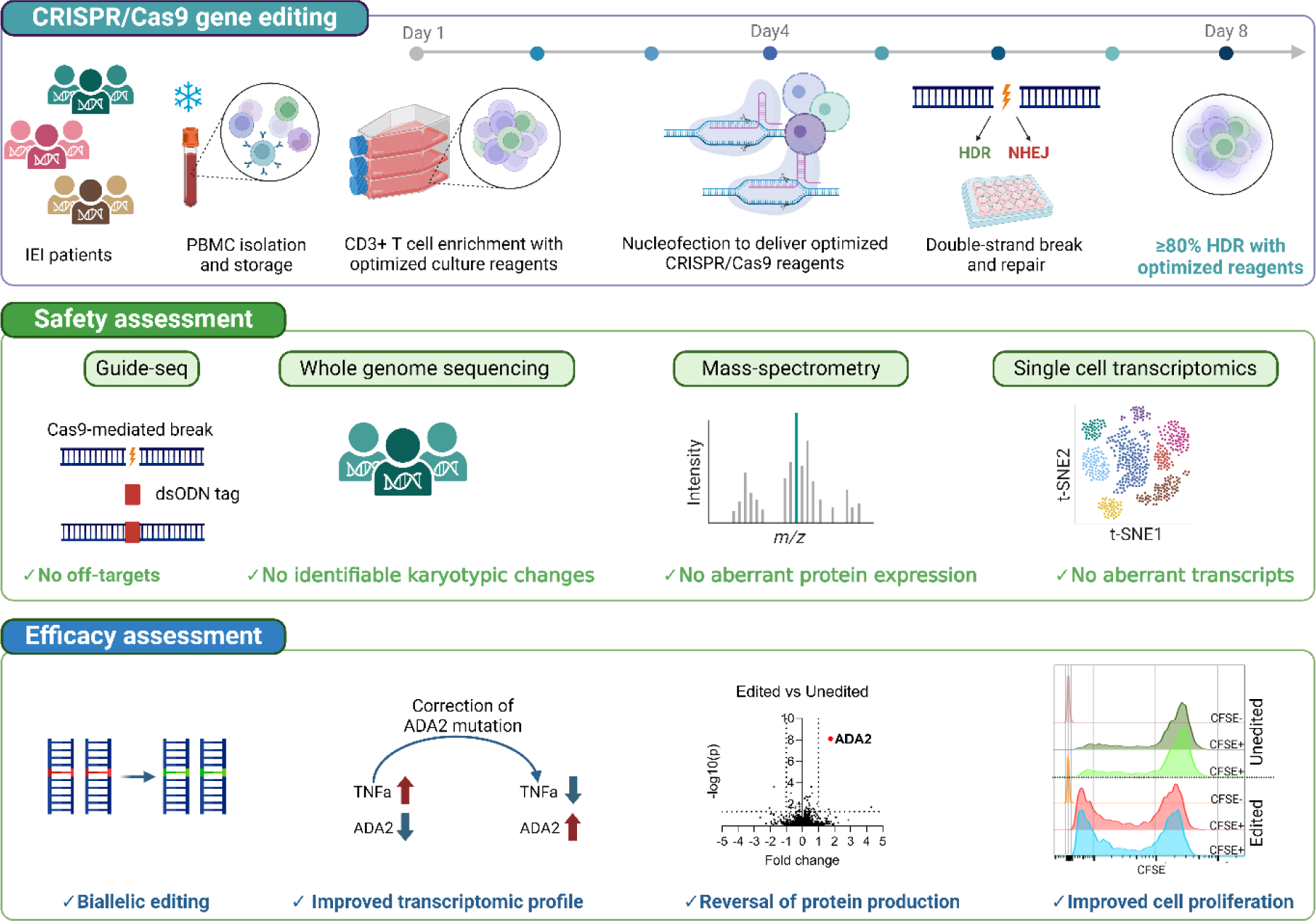
GRAPHICAL ABSTRACT.

## INTRODUCTION

Inborn errors of immunity (IEI) encompass ∼500 single-gene defects that lead to diverse clinical presentations, including infection susceptibility, autoimmunity and –inflammation, cancer predisposition, and allergies[1, 2]. IEIs are popular targets for CRISPR-Cas9 gene correction. For some IEIs, knockout of the disease gene can restore normal cell function[3, 4]. However, the main correction strategy is the knock-in of the healthy cDNA under the endogenous promoter of the diseased gene, or into the safe harbor locus[5, 6]. The strategy can treat most of the defects caused by the given single gene, but sometimes results in disordered gene expression and is slow to adapt to large genes, novel disease gene discoveries or ultra rare IEIs, which might feature only <10 patients globally.

Precise correction of the pathogenic variant can circumvent the problems of therapeutic cDNA knock-in. However, the cost-effective clinical translation of precise gene correction would require a platform approach where many IEIs can be treated with a streamlined protocol where only the guide and the repair template change. Precise correction of monogenic mutations is typically attributed to base[7, 8] and prime editing[8, 9], as these methods do not induce double-strand breaks and are thus considered the safest alternative. However, guide design options for base editors are often limited, with 1-3 available guides per target and a risk for bystander editing that changes the coding bases nearby. Prime editing guide RNA (pegRNA) design for prime editing is even more challenging, as the rules for effective pegRNA design are incompletely understood, and a large screen is often necessary to identify an effective pegRNA for a given patient mutation[10]. The guide and repair design rules are better defined for the “standard” CRISPR-Cas9 –driven homology-directed repair, and the strategy can target virtually all SNVs and small indels in the human genome. The main drawback has been the method safety, as well as the requirement of the S phase of the cell cycle (and the resulting need for cell division) to complete the repair.

In this study, we have developed a single nucleotide variant (SNV) T cell editing protocol that utilises CRISPR-Cas9 –driven homology-directed repair (HDR) and can correct mutations with up to 80% efficiency. Hematopoietic stem cell editing is considered the prime target for IEI correction; however, in IEIs with faulty T cell function, therapeutic benefit can also be achieved by correcting patient T cells. The cells can then be infused to the patient as an auto transplant to control viral infections and other pathology that stems from faulty T cell function[11]. T cell therapy would particularly benefit patients who cannot tolerate stem cell transplant – allogeneic, or with autologous corrected stem cells – due to their advanced disease, acute infections, or lack of a suitable donor[11, 12]. Our study uses the following Finnish founder diseases as models, as we had sizeable cohorts of homozygous patients available for study: ADA2 deficiency (*ADA2* rs77563738, p.R169Q), Autoimmune polyendocrinopathy-candidiasis-ectodermal dystrophy (APECED) (*AIRE* rs121434254, p.R257X) and Cartilage Hair Hypoplasia (CHH) (*RMRP* rs199476103, c.A71G). ADA2 p.R169Q has no available base editing guides, and the AIRE and RMRP guides can induce bystander base editing (Supplementary Figure 1), rendering comparisons with other editing methods difficult.

During protocol development, we investigated several novel and published strategies for enhancing homology-directed repair and improving T cell expansion and viability. We show that guide RNA selection is crucial for maximum efficiency and safety and present an improved protocol for performing GUIDE-seq off-target profiling in patient T cells. Finally, we screened the genomic integrity of the edited cells with long read whole genome sequencing, single cell RNA sequencing, and proteomic studies, and did not detect pre-malignant karyotypic, transcriptomic and proteomic aberrations with our chosen guides. Our pipeline is non-viral and compatible with GMP scale-up. It also edits CD34+ hematopoietic stem cells with up to 80% efficiency, suggesting that the presented optimizations have the potential to extend to other primary cell types.

## MATERIALS AND METHODS

### Isolation, culture, and nucleofection of T cells, CD34+ HSPCs, and fibroblasts

We obtained peripheral blood, cord blood, and skin biopsies from patients or healthy donors. Detailed information on the patients is provided in Supplementary Table 1. The study was conducted in accordance with the principles of the Helsinki Declaration and was approved by the Helsinki University Central Hospital Ethics Committee, and the Regional Committee for Medical and Health Research Ethics South-East Norway. Participants have signed written informed consent.

PBMCs were isolated from peripheral blood using Ficoll gradient centrifugation and then cryopreserved. Upon thawing, PBMCs were cultured in ImmunoCult™-XF T Cell Expansion Medium supplemented with IL-2, IL-7, IL-15, and CD3/CD28 T Cell Activator. After 3 nights at 37°C/5% CO_2_, cells were nucleofected or further cultured without CD3/CD28 Activator. CD34+ HSPCs were isolated from cord blood using CD34 MicroBead Kit and then cryopreserved. Upon thawing, CD34+ cells were cultured in StemSpan™ SFEM II supplemented with GlutaMax, Flt3-L, TPO, SCF, IL-6, StemRegenin-1 and UM729. After 3 nights at 37°C/5% CO_2_, cells were nucleofected or further cultured. Fibroblasts isolated from skin biopsies were expanded in DMEM with low glucose, pyruvate, and FBS, and cryopreserved. Upon thawing, fibroblasts were cultured until confluent, passaged every 3-4 days with TrypLE™ Express Enzyme, and nucleofected by passage 10.

T cells, CD34+ HSPCs, and fibroblasts were nucleofected using a 4-D Nucleofector system and 96-well unit (Lonza). gRNAs were prepared by annealing crRNA and trcrRNA (IDT) and mixed with Cas9 nuclease and ssODN (IDT) to form RNPs. T cells (0.5 or 1 million), HSPCs (0.3 million), and fibroblasts (1 million) were resuspended in 20 µL electroporation buffer and nucleofected using programs EO-115, DZ-100, and CA-137, respectively. Post-nucleofection, T cells were incubated with recovery medium (basal medium supplemented with IL-2) for 15 minutes, transferred to plates, and cultured until collected after 4-8 days. HSPCs were incubated for 15 minutes with a stimulation medium (basal medium supplemented with aforementioned cytokines), transferred to plates and cultured until collected after 4 days. Fibroblasts were incubated with culture medium (basal medium supplemented with aforementioned cytokines) for 15 minutes, transferred to plates, and cultured until collected after 4 days. Detailed descriptions of the methods can be found in Supplementary Methods.

### Design and screening CRISPR/Cas9 reagents

Seven to eighteen gRNAs were designed based on available PAM (NGG) sites within the 100 bp repair template region centering the mutation site. Single-stranded DNA repair templates (ssODNs) of 100 bp were designed with ±50 bp homology arms. Synonymous, silent SNVs were added in repair templates for ADA2 (four SNVs) and AIRE (three SNVs) to prevent CRISPR re-cutting and ensure identical editing in donors and patients. As RMRP is a non-coding gene, SNVs were used in early experiments, and only mutation correction was used for functional assessments. Asymmetric ssODNs with 10-40 bp homology arms were tested to enhance homology-directed repair. Details related to gRNA and ssODN design, BG-coupled repair template oligos and Cas9-SNAP protein production can be found in Supplementary Methods.

### On-target editing assessment

ddPCR assays were conducted to assess HDR and NHEJ editing. Previously described oligos[13] were used for Enh4-1, CTCF1, and RNF2, while new oligos for ADA2, AIRE, and RMRP were designed. ddPCR was performed using the QX200 system (Bio-Rad) and analyzed with QuantaSoft software (Bio-Rad). The oligos are listed in Supplementary Methods.

Amplicon sequencing libraries were prepared from gDNA samples using a two-step PCR method[13]. Unique Molecular Identifiers were added to the primers to filter out PCR bias[13]. Libraries were sequenced using the Illumina MiSeq v2 platform. Data analysis was done using the ampliCan software package[14].

### Assessment of *in silico* gRNA design tools

We assessed the predictive power of *in silico* gRNA design tools against *in vitro* gRNA screening data using the following tools: Atum, Benchling, CHOPCHOP, CRISPOR, DeepSpCas9, EuPaGDT, and IDT gRNA design tool. Using 100 bp mutant-specific sequences with 50 bp homology arms as input, we selected three highest predicted efficiency gRNAs from each tool. These were then compared against the three best *in vitro*-validated gRNAs from patient T cells. Details of *in silico* tools are listed in Supplementary Methods.

### Screening HDR enhancers and cell cycle inhibitors in healthy donor T cells

33 HDR-enhancers and 10 cell cycle inhibitors (described in Supplementary Methods Table 10) were screened in HD T cells at three concentrations against a DMSO-treated RNP-edited cells. For HDR enhancers, 0.5 million T cells per sample were nucleofected and incubated with the compounds in T cell recovery medium for 24h, then split 1:1 in recovery medium without compounds at 24h and 72h. Toxicity of HDR-enhancers was assessed using the CellTiter-Glo assay (Promega) according to the manufacturer’s instructions. The detailed protocol can be found in Supplementary Methods. For cell cycle inhibitors, cells were either pre-treated with the compounds for 24h before nucleofection or 24h after nucleofection. In both cases, 0.5 million cells per sample were nucleofected and split 1:1 in recovery medium without compounds at 24h and 72h. Samples for both screens were collected for gDNA extraction and ddPCR 96h after nucleofection.

### Off-target editing assessment

The previously published Genome-wide, unbiased identification of DSBs enabled by sequencing (GUIDE-seq)[15] was used to assess the off-target editing. The detailed protocol is described in Supplementary Methods. In brief, 1 million T cells per sample were nucleofected on day 5 with RNPs (100 pmol gRNAs, 61 pmol Cas9, 30 pmol dsODN). After nucleofection, cells were transferred to 24-well plates with 500 µL recovery medium and split at 24h and 72h. Samples were collected for library preparation, sequencing and ddPCR 4 days later. Data analysis was performed following the GUIDE-Seq analysis pipeline from Zhu et al. 2017[16], but adjusted for allowing bulges between sgRNA and off-target sites with editing distance of 4. Final off-targets were normalized against control data (transfected with dsODN only). We used custom scripts available at https://git.app.uib.no/valenlab/t_cell_editing_pipeline/.

### PacBio sequencing of CRISPR-edited healthy donor T cells

Healthy donor T cells were edited as described above, 0.5 μM KU0060648 or DMSO. Six days post-editing, DNA was extracted from 5 million cells per sample using Qiagen kits. DNA quality was assessed using NanoDrop, Qubit, and agarose gel electrophoresis. Libraries for PacBio HiFi sequencing were prepared using the Revio HiFi prep kit and Sequencing chemistry v2.0. Sequencing data was demultiplexed with SMRT Link, and CCS reads were generated and further demultiplexed using barcoded primers, with HiFi reads indexed by barcode IDs. The HiFi sequencing reads were aligned with pbmm2 v1.13.0. Structural variants were called with pbsv v2.9.0, small variants with deepVariant v1.6.0. All possible mismatches, deletions and insertions were extracted from aligned reads using custom scripts (https://git.app.uib.no/valenlab/t_cell_editing_pipeline/-/tree/main/katariina_pacbio). We normalized data using two control samples and focused on sites that were potential sgRNA off-target within distance of 4, allowing for bulges.

### Immunophenotyping by Flow cytometry

PBMC samples from days 1, 4, and 8 of the editing pipeline were assessed using flow cytometry. Cells (0.5 million per sample) were washed with flow cytometry buffer, blocked with 10% human serum, and stained with an antibody cocktail (described in Table 3 in Supplementary Methods) to identify CD4 T cells, CD8 T cells, B cells, NK cells, monocytes, and dendritic cells. After washing, cells were resuspended in 250 μL flow cytometry buffer and stored at 4°C. Flow cytometry was done on LSRII and data analysis was done using FlowJo. For detailed protocol see Supplementary methods.

### T cell proliferation assay in CHH patients

T cells from CHH patients from day 20 of the editing pipeline were collected, washed with PBS, and resuspended at 2 million cells/mL. Cells were stained with 1 μM CFSE and incubated in the dark at 37°C for 5 minutes. Cold human serum was added to quench the reaction. The cells were then washed and resuspended in Immunocult medium supplemented with 250 U/mL IL-2 and 0.2 million cells were plated in 96-well U-bottom plate. After four days, cells were stained with an antibody cocktail (described in Table 5 in Supplementary Methods) and analyzed by flow cytometry as described in the previous section (Immunophenotyping by flow cytometry). For detailed protocol see Supplementary methods.

### DNRT Single-cell RNA sequencing and quantitative real-time PCR of control and DADA2 patient T cells

Previously published, Smart-Seq2 –based DNTR (Direct Nuclear Tagmentation and RNA sequencing) protocol was used[17]. For detailed protocol see Supplementrary methods. In brief, on day 8, nucleofected T cells from DADA2 patients and healthy donors were collected, washed, and stained with Live/Dead dye and Fc blocking reagent. After washing, the cells were stained with antibody cocktail (described in Table 4 in Supplementary Methods). The cells were then washed and resuspended in flow buffer before sorting live CD4+ and CD8+ T cells into 384-well plates with lysis buffer. Post sorting the plates were centrifuged, snap-frozen, and stored at –80°C. Using the Smart-Seq2 protocol[17], cells were thawed, reverse transcribed, and cDNA pre-amplified, with cleanup using SPRI beads and concentration measured with Qubit DNA HS kit. Tagmentation of diluted cDNA was followed by SDS reaction stop, barcoding, and PCR. Libraries were cleaned with SPRI beads and sequenced on a Novaseq 6000.

For data analysis, the reads were trimmed with Cutadapt[18] and aligned to hg38 with STAR[19]. Picard[20] removed duplicates, and HTSeq[21] summarized counts. Cells with <20,000 reads, <500 features, or low ACTB expression were filtered out. Seurat[22] version 5.0.1 log-normalized data, identified 2000 variable features, and scaled data per condition. FindMarkers in Seurat identified markers between conditions, and fgsea[23] performed gene set enrichment analysis. Fusion gene detection was performed with STAR-fusion, see supplementary methods. LOH calculations were performed as described in Supplementary methods. For quantitative analysis of different alleles in single cells, 1 µL of diluted cDNA was amplified with specific probes for WT and edited alleles. Detailed protocol can be found in Supplementary Methods. qPCR analysis used BioRad software with a 200 RFU as a threshold for determining which allele was being expressed.

### Mass spectrometry

For details see Supplementary methods. In brief, T cells from three DADA2 patients and healthy donors were cultured with 1 million cells per sample and nucleofected on day 5. Mock –nucleofected cells were treated with DMSO, and edited cells were treated with 0.5 μM KU0060648, 0.6 μM IDT Alt-R enhancer V2, or DMSO for 24h. Cells were collected on day 12, washed, pelleted, and snap-frozen on liquid nitrogen.

For mass spectrometry, Trypsin/LysC digested samples were diluted 1:60 in 0.1%FA in water, and 20µL was loaded into an Evotip. Samples were analyzed using an Evosep One system with a Bruker timsTOF Pro mass spectrometer. Peptide separation used an 8 cm × 150 µm column with a 21 min gradient. Data was processed with DIA-NN v1.8.1[24, 25]using the UniProt human proteome spectral library, with fixed and variable modifications. Pre-processing involved log2 transformation, median-normalization, and QRILC imputation (Lazar & Burger, 2022, available at: https://cran.r-project.org/web/packages/imputeLCMD/imputeLCMD.pdf). Statistical analysis used student’s t-test[26]and the Benjamini-Hochberg method[27] for p-value adjustment. The volcano plots were generated using bioinfokit.

### Data availability

The manuscript contains the following supplementary tables (Excel unless otherwise noted):

- Supplementary Methods (PDF)
- Supplementary Figures (PDF)
- Supplementary Tables 1-6 (individual Excel files)

Raw GUIDE-seq, scRNA-sequencing, mass spectrometry and WGS data will be deposited in a secure repository after publication.

## RESULTS

### Repair strategy

Guide design is crucial for the success of CRISPR experiments[28]. To identify gRNAs that incite high templated correction in the three model loci, we screened all the available guides that cut within a 100 bp repair template region (7-18 guides per locus). To prevent CRISPR re-cutting[29, 30], enable identical ssODNs for healthy and patient cells, and allow rapid detection of homology-directed repair by droplet digital PCR (ddPCR), we designed a repair strategy where 3-4 silent SNVs were added close to the mutation site in addition to mutation correction (Figure 1A-B, Supplementary Figure 1A,E,I). Since RMRP encodes a non-coding RNA, we could not design silent SNVs to the locus and thus knocked in two variants of unknown function during the early optimization experiments (Supplementary Figure 1I-J). In later studies, we only corrected the pathogenic variant (Supplementary Figure 1K).

**Figure 1.**
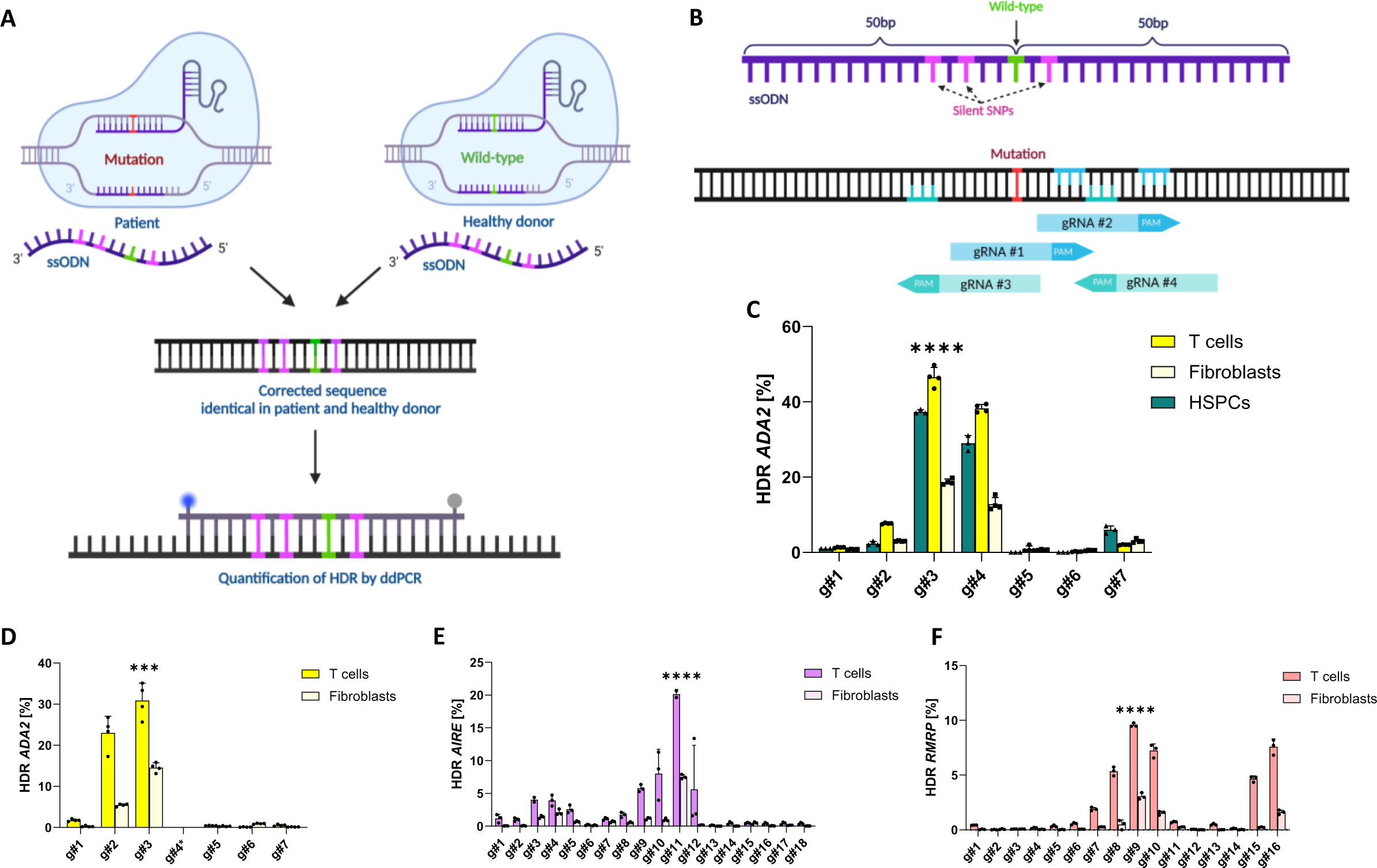
Repair template design and gRNA screening in patient T cells and fibroblasts. (**a**) Schematic representation of the editing strategy used in the study for ADA2 and AIRE. In short, 100bp ssODNs with +/-50bp homology arms to the mutant site were designed for editing patient and healthy control cells, where in addition to correcting the mutation (mutation marked in red, correction in green), 3-4 silent SNPs (pink) were added into the repair template. To study HDR editing in cells, ddPCR was used with probes specifically binding to the HDR edited site. As RMRP is non-coding, non-silent SNPs were used in early experiments and mutation correction later for functional assessments. **(b)** Schematic representation of the gRNA design used in the study. In short, multiple gRNAs per locus were designed based on available PAM sites surrounding the mutation site (red). Forward gRNAs and their respective PAM sites are marked in blue and reverse gRNAs and their PAMs in turquoise. **(c)** ADA2 gRNA screening in HD T cells, fibroblasts and CD34+ HSPCs, assessed by ddPCR (measurements performed in quadruplicates in T cells and fibroblasts and triplicates in HSPCs). **(d)** ADA2 gRNA screening in DADA2 patient T cells and fibroblasts, assessed by ddPCR (measurements performed in triplicates). ADA2 g4, marked with asterisk, was not tested in patients due to loss of PAM site caused by the mutation. **(e)** AIRE gRNA screening in APECED patient T cells and fibroblasts, assessed by ddPCR (measurements performed in triplicates). **(f)** RMRP gRNA screening in CHH patient T cells and fibroblasts, assessed by ddPCR (measurements performed in triplicates). One independent experiment was performed for all sets of data. Statistical significance of highest HDR for a given gRNA was assessed by one-way ANOVA with Fisher’s LSD test, where ****p<0.0001 and ***p<0.0002. Bar denotes mean value, error bars represent ± SD. Abbreviations: HD (healthy donor), HDR (homology-directed repair), ddPCR (Droplet Digital PCR), gRNA (guide-RNA), ssODN (single-stranded oligodinucleotide), PAM (protospacer adjacent motif), nt (nucleotide), DADA2 (Deficiency of adenosine deaminase 2), APECED (Autoimmune polyendocrinopathy-candidiasis-ectodermal dystrophy), CHH (Cartilage hair hypoplasia).

We first tested the correction efficiency for the ADA2 locus in healthy control T cells, fibroblasts, and cord blood hematopoietic stem cells and compared the results to similar screens in DADA2 patient T cells and fibroblasts (Figure 1C-D; all patients are homozygous for the ADA2 p.R169Q mutation). The best guide outperformed the suboptimal ones regardless of the cell type, indicating flexibility in using different patient primary cell types for initial guide screening. We identified gRNA #3 as the best guide for ADA2 correction, with ∼30% maximum HDR efficiency (Figure 1D). We then screened guides for AIRE and RMRP loci in homozygous patient T cells and fibroblasts and identified AIRE gRNA #11 and RMRP gRNA #9 as the best guides (Figure 1E-F, 10-20% templated correction). We assessed the editing by ddPCR and deep amplicon sequencing with near identical results (Supplementary Figure 1C,G,L), confirming ddPCR as a reliable method for rapid HDR assessment. However, *in silico* gRNA design tools showed poor accuracy with this design strategy, likely as they are built on datasets adapted for NHEJ-based gene knockout (Supplementary Figure 1D,H,M).

### Optimized T cell culture for editing enhancement

HDR-dependent correction happens in the S/G2 cell cycle phase[31]. Therefore, optimal T cell expansion and viability can further increase gene correction[32]. As a baseline, we used common T cell editing protocols and our previous work[13, 33, 34] where Peripheral Blood Mononuclear Cells (PBMCs) are first stimulated for three days, then nucleofected with CRISPR-Cas9 ribonucleoprotein complexes (RNPs), and collected for DNA extraction 3-5 days post electroporation (Figure 2A). We noted ∼5-8% AIRE and ∼15-30% ADA2 HDR editing in healthy donors, with editing levels plateauing two (AIRE) and three (ADA2) days after nucleofection and staying consistent for up to 14 days after nucleofection (Supplementary Figure 2A-B). The cell quantity had no effect on the final editing level, allowing us to work with less material when necessary (Figure 2B). We thus settled for 0.5M-1M cells per nucleofection to allow enough material for downstream analyses and standardized the sample collection on day four post-nucleofection.

**Figure 2.**
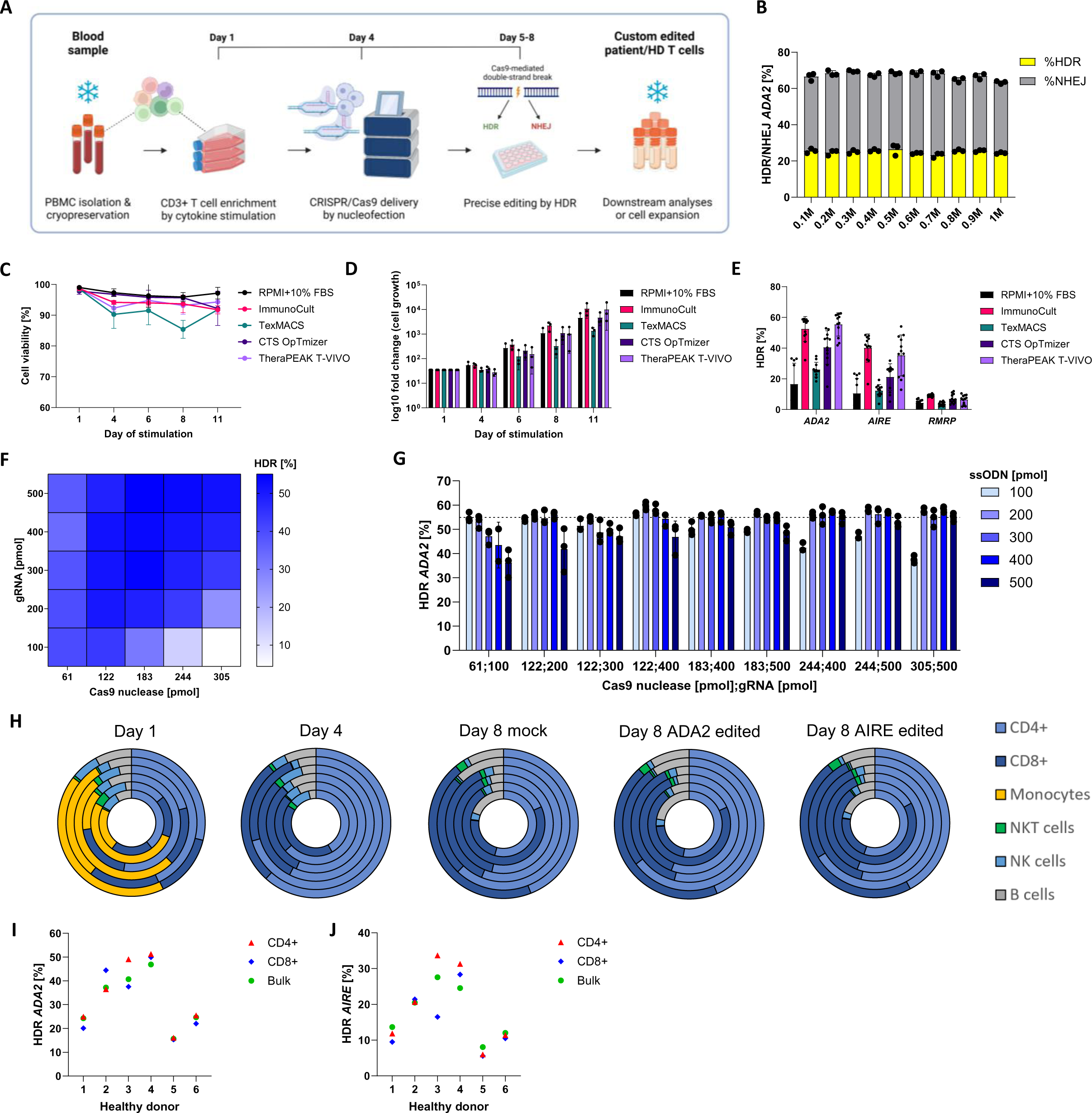
Establishment and assessment of CRISPR/Cas9 T cell editing pipeline. (**a**) Schematic representation of the 8-day CRISPR/Cas9 T cell pipeline. The workflow starts with isolation of PBMCs from patient or HD blood samples. PBMCs can be cryopreserved and thawed, or cultured fresh to initiate the pipeline, briefly discussed here: on day 1, PBMCs are stimulated for three days with interleukins: IL-2 (120 U/mL), IL-7 (3 ng/μL), IL-15 (3 ng/μL) and soluble CD3/CD28 (15 μL/mL), which activate and induce expansion of CD3+ T cells. Cells are nucleofected on day 4 to deliver CRISPR reagents (gRNA, Cas9 nuclease, ssODN) into nuclei of the cells for genomic editing. After nucleofection, cells are cultured for four days in IL-2 (250 U/mL), during which cells repair the Cas9-mediated double stranded break by HDR or NHEJ. Cells are harvested on day 8 for downstream assays, cryopreserved for later use or expanded further. **(b)** ADA2 HDR and NHEJ editing in HD T cells with 0.1-1M nucleofected cells/sample, measured by ddPCR (measurements performed in triplicates). Comparison of different T cell culture medium during 11-day cytokine stimulation, assessed by **(c)** T cell viability (each dot represents the mean measurement from three HDs, n=3 experiments), **(d)** T cell fold change (each dot representing one HD, n=3 experiments) and **(e)** HDR editing on day 8 for ADA2, AIRE and RMRP (one dot represents one technical measurement out of triplicate measurements per condition, n=3 experiments). **(f)** ADA2 HDR editing in HD T cells with Cas9 nuclease at 61-305pmol/sample, gRNA at 100-500pmol/sample and ssODN at 100pmol/sample, measured by ddPCR (measurements performed in triplicates). **(g)** ADA2 HDR editing in HD T cells with selected combinations of RNP concentrations with ssODN at 100-500pmol/sample, measured by ddPCR (measurements performed in triplicates). Dashed line indicates mean measurement for Cas9 nuclease at 61pmol, gRNA at 100pmol and ssODN at 100pmol/sample. **(h)** Frequency of immune cells (CD4+, CD8+, monocytes, NKT cells, NK cells, B cells) in six HDs on day 1, 4 and 8 (mock, ADA2 or AIRE edited) of the pipeline, assessed by flow cytometry. Each ring of the doughnut plot represents one donor. HDR editing levels for ADA2 **(i)** and AIRE **(j)** in six HDs for CD4+, CD8+ and bulk of cells on day 8, measured by ddPCR (measurements performed in single replicas). One independent experiment was performed for all sets of data except for (c)-(e) where data from three donors are shown in the graphs and (f)-(g) where one out of three representative experiments is shown. Bar denotes mean value, error bars represent ± SD. Abbreviations: PBMC (peripheral blood mononuclear cell), HD (healthy donor), IL (interleukin), gRNA (guide-RNA), ssODN (single-stranded oligodinucleotide), HDR (homology-directed repair), ddPCR (Droplet Digital PCR), RNP (ribonucleoprotein).

To improve T cell proliferation, viability and HDR, we compared several GMP-compatible cell media while editing ADA2, AIRE and RMRP loci (Figure 2C-E). Based on the results across the tested donors, Immunocult and TheraPEAK T-VIVO performed similarly but as Immunocult supported cell proliferation earlier in the pipeline, we selected it, combined with 120 U/mL IL-2, 3 ng/μL IL-7, 3 ng/μL IL-15 and 15 μL/mL soluble CD3/CD28. We also titrated the concentrations of Cas9 nuclease, gRNA, and repair templates, with the goal of reaching optimal reagent concentration in the nucleus without excessive toxicity (Figure 2F-G, Supplementary Figure 2C-D). Based on the results, we standardised Cas9 nuclease at 61pmol, gRNA at 100pmol, and ssODN at 100pmol per sample.

To verify selective CD3+ T cell expansion from PBMCs, we quantified the immune cell populations on culture days 1, 4, and 8 from six healthy donors by flow cytometry (Figure 2H). While PBMC population diversity is considerable on day 1, it gradually disappears during cytokine stimulation. By day 8, CD4+ and CD8+ T cells make up ∼80% of all cells. On day 8, it is possible to sort T cells, cryopreserve for later use or expand further to obtain a pure T cell population. Although we noted significant interindividual and locus-specific variation in editing efficiency as also described by others[13, 32, 35, 36], HDR editing levels in CD4+ and CD8+ subsets were similar within a donor. (Figure 2I-J).

### Refined repair template design for editing enhancement

The positioning and format of the repair template affects HDR editing[30, 37–39]. For optimal template positioning, we designed asymmetric 100 bp templates for ADA2, AIRE, and RMRP loci, with 10-90bp homology arms on either side (9 templates per locus, Figure 3A)[37]. The symmetric templates with 50bp homology arms proved best for ADA2 and RMRP; however, AIRE locus edited most optimally with an asymmetric template (30bp left, 70pb right homology arm, Figure 3B-D).

**Figure 3.**
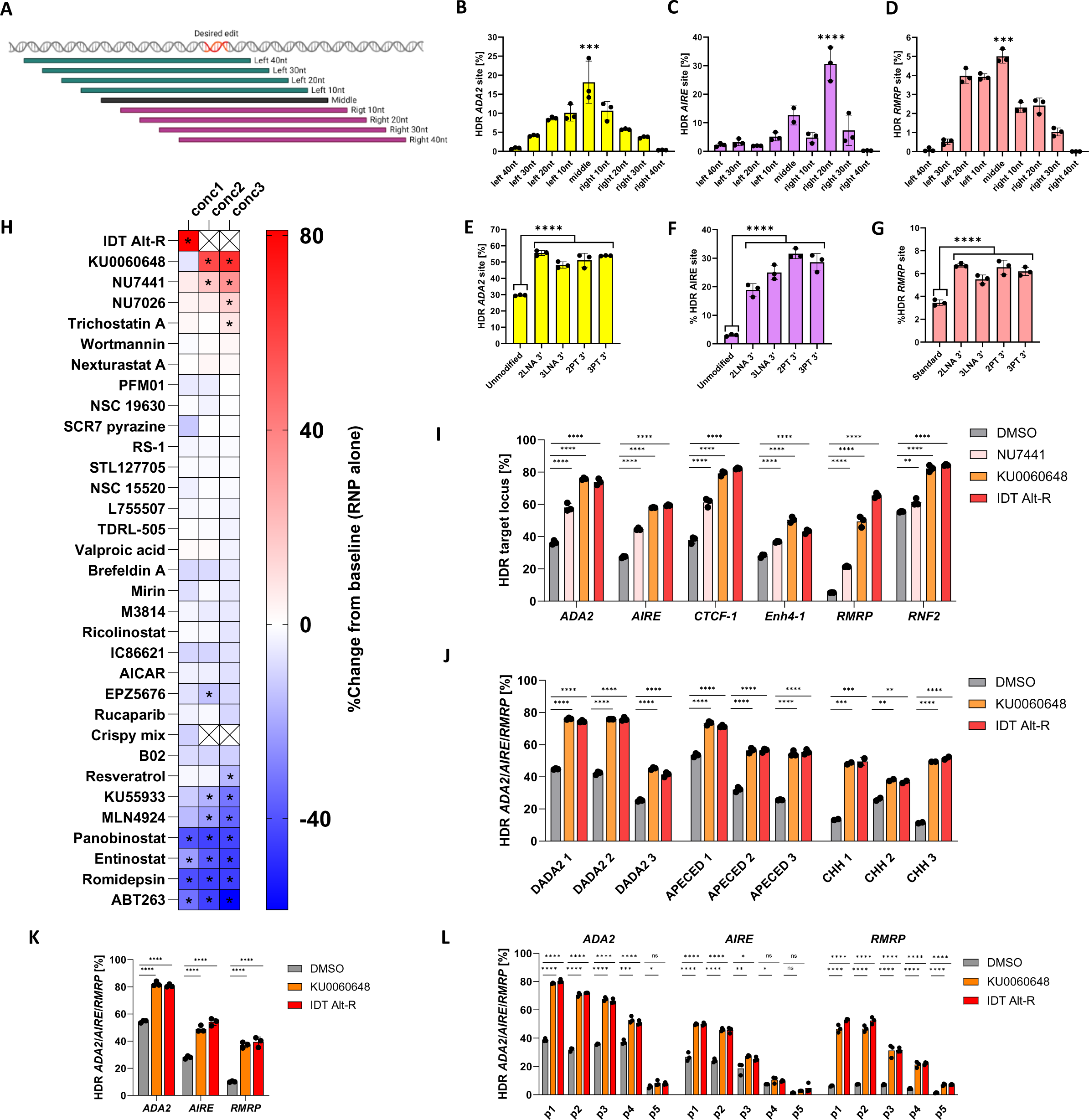
Improvement of HDR editing in primary HD and patient T cells. (**a**) Schematic representation of asymmetric ssODN designs with 10-90bp homology arms on either side. Quantification of HDR editing with asymmetric ssODNs in HD T cells for **(b)** ADA2, **(c)** AIRE and **(d)** RMRP, measured by ddPCR (measurements performed in triplicates). Quantification of HDR editing with 3’ LNA-or 3’PT-modified ssODNs with position-optimized ssODNs in HD T cells for **(e)** ADA2, **(f)** AIRE and **(g)** RMRP, measured by ddPCR (measurements performed in triplicates). **(h)** Validation of HDR enhancing compounds at three concentrations (conc1-3) for ADA2 in HD T cells, assessed by ddPCR (measurements performed in duplicate). Compounds were tested in three HDs in duplicate measurements per condition, where mean value of the measurements from all donors per condition was compared to mean value of DMSO baseline (ADA2 RNP alone). Statistical significance of compounds was assessed by ANOVA. For the heatmap, percentage of HDR fold change from baseline was calculated for each concentration. Statistically significant concentrations are indicated in black asterisks. Conc2-3 are marked with a cross for compounds that were only assessed at one concentration. **(i)** Quantification of HDR editing for ADA2, AIRE, CTCF1, Enh4-1, RMRP and RNF2, measured by ddPCR (measurements performed in triplicates) with selected HDR enhancing small molecules (4 μM NU7441, 0.5 μM KU0060648, 1 μM IDT Alt-R enhancer V2) or DMSO (RNP alone) in HD T cells. **(j)** Quantification of HDR editing for mutation correction in DADA2, APECED and CHH patients with concentration-optimized HDR enhancing small molecules (0.5 μM KU0060648, 0.6 μM IDT Alt-R enhancer V2) where ADA2, AIRE and RMRP loci, respectively, were corrected. HDR levels were assessed by ddPCR for DADA2 and APECED patients (measurements performed in triplicates) and by amplicon sequencing for CHH patients (measurements performed in duplicates). **(k)** Quantification of ADA2, AIRE and RMRP HDR editing in HD CD34+ HSPCs, measured by ddPCR (measurements performed in triplicates) with concentration-optimized HDR enhancing small molecules (0.5 μM KU0060648, 0.6 μM IDT Alt-R enhancer V2) or DMSO (RNP alone). **(l)** Quantification of HDR editing for ADA2, AIRE and RMRP in HD T cells at different cell passages (p1-p5), measured by ddPCR (measurements performed in triplicates) with concentration-optimized HDR enhancing small molecules (0.5 μM KU0060648, 0.6 μM IDT Alt-R enhancer V2) or DMSO (RNP alone). Three independent experiments were performed for all sets of data where representative experiment is shown except for (h) where average measurements from three healthy donors is shown, (j) where all three patients are shown in the graph and (k) where compounds were tested in one donor. Bar denotes mean value, error bars represent ± SD. Statistical significance for all sets of data, except (h), was assessed by one-way ANOVA with Fisher’s LSD test, where ****p<0.0001, ***p<0.0002, **p<0.001 and *p<0.01. Abbreviations: HD (healthy donor), ssODN (single-stranded oligodinucleotide), nt (nucleotide), HDR (homology-directed repair), ddPCR (Droplet Digital PCR), RNP (ribonucleoprotein), LNA (locked nucleic acid), PT (phosphorothioate), DADA2 (Deficiency of adenosine deaminase 2), APECED (Autoimmune polyendocrinopathy-candidiasis-ectodermal dystrophy), CHH (Cartilage hair hypoplasia), HSPC (hematopoietic stem and progenitor cell.

Coupling the repair template to Cas9 has been shown to improve HDR editing, presumably by enhancing nuclear import and template positioning at the cut site[40, 41]. To test the strategy, we synthesised 5’ benzylguanine (BG)-coupled repair templates that can covalently link to Cas9-SNAP fusions that target the ADA2 locus[41, 42]. BG-templates led to two-fold editing enhancement with both Cas9WT and Cas9-SNAP RNPs, in fibroblasts and T cells (Supplementary Figure 3A-C). We concluded that the observed enhancement is likely explained by the BG modification stabilizing the template, with Cas9 coupling being secondary to this effect. However, we did not pursue this approach due to difficulty in assessing the purity of the in-house produced BG oligos. Nevertheless, knowing that the modification of repair template can increase HDR efficiency, we then tested different phosphorothioate (PT)– and locked nucleic acid (LNA) modifications in different nucleotide positions that stabilize the repair template and were easily available commercially (Supplementary Figure 3D-F). 3’ PT and LNA modifications led to two-fold HDR improvements (Figure 3E-G). Therefore, we chose 2PT 3’ modification of the repair templates for further experiments due to its universal effectiveness and ease of synthesis (Supplementary Figure 3G-I).

### Inhibition of DNA-dependent protein kinase further improves homology-directed repair

A considerable number of HDR-enhancing chemicals have been published. To see whether we could further improve editing efficiency, we selected 33 chemical compounds convincingly reported as HDR enhancers (Table 10 in Supplementary Methods) and screened them for editing enhancement in the ADA2 locus in healthy donor T cells. Most of the reported compounds decreased HDR, likely due to cell toxicity. Three compounds showed two-fold improvement, leading up to 80% efficiencies in screening conditions (Figure 3H): DNA-dependent protein kinase (DNA-PK) inhibitors NU7441[43] and KU0060648[43], and IDT Alt-R enhancer V2 (hereby referred to as IDT Alt-R). We validated these three compounds in six endogenous loci, optimized their concentrations in healthy donors and tested them further in DADA2, APECED, and CHH patient T cells, consistently achieving minimal toxicity, ∼two-fold improvement and up to 80% mutation correction, depending on the target locus and individual (Figure 3I-J, Supplementary Figure 3J-K). Furthermore, the compounds improved editing even in cord blood hematopoietic stem cells (Figure 3K). High HDR levels were maintained when nucleofection was performed at passages 1-3, decreasing in later passages (Figure 3L).

As HDR is dependent on the S/G2 phase of the cell cycle, we also tested a set of cell cycle inhibitors (Table 11 in Supplementary Methods) for their ability to synchronize editing to S/G2 phase and consequently increase HDR. Hydroxyurea[44] emerged as an unexpected editing enhancer when applied 24h before nucleofection, but as the effect was suboptimal in comparison to NHEJ inhibition, we did not explore the strategy further (Supplementary Figure 3L-M).

### Adapted GUIDE-seq off-target profiling for patients and healthy donors

GUIDE-seq finds CRISPR off-target cutting by transfecting cells with modified double-stranded DNA oligos (dsODNs) along the CRISPR RNP complex, and then selectively amplifying and sequencing the oligo integration sites[15]. The existing GUIDE-seq data mainly comes from cell lines[15]. There are reports for adaptations to T cells[33, 45], but since dsODN can be toxic particularly to patient T cells, we started the off-target profiling by optimising the dsODN concentration for improved cell viability. Experiments in healthy donor T cells for guides targeting the ADA2 and HEK-site 4 loci (positive control guide with multiple off-targets[15]) showed acceptable cell viabilities and optimal dsODN integration at 20-100 pmol dsODN per sample. Subsequent deep sequencing detected no off-targets for ADA2 guide but recovered several integrants for HEK-site 4, validating the method’s sensitivity (Supplementary Figure 4A-F).

To account for increased dsODN toxicity in patient T cells, we refined the dsODN concentration further in DADA2 patient T cells, settling on 30 pmol/sample based on cell viability, dsODN integration and cell yield (Figure 4A-C). Finally, we performed GUIDE-seq in three patients and three healthy donors for each locus and confirmed the safety of ADA2 gRNA #3, AIRE gRNA #11 and RMRP gRNA #9 with no off-targets, contrasting with multiple off-targets for HEK-site 4 (Figure 4D-H). To summarise, we present a refined protocol for T cell CRISPR off-target profiling and recommend lower dsODN concentrations for IEI patient samples to reach optimal cell viability and reliable sequencing results.

**Figure 4.**
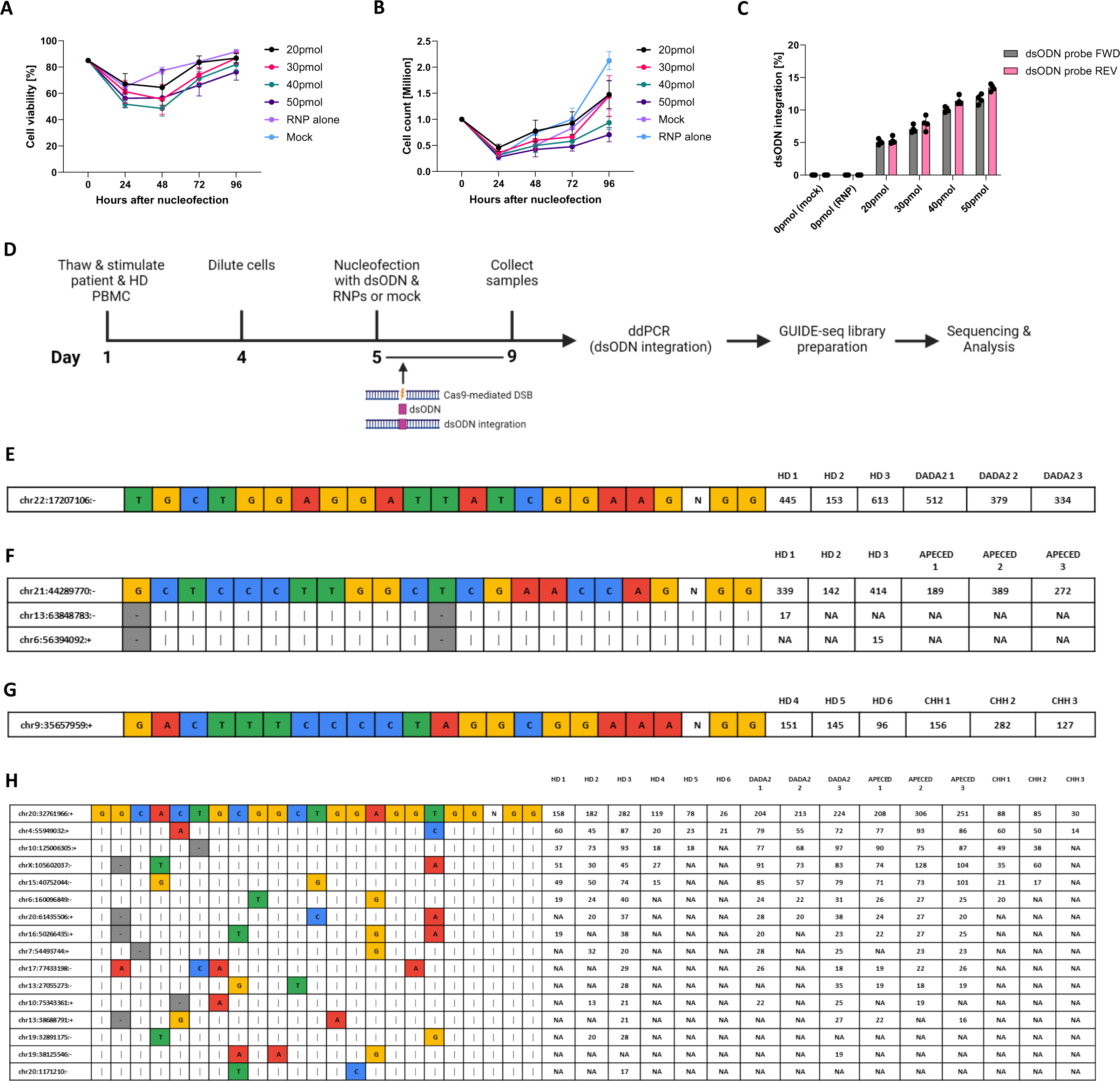
Guide-RNA off-target profiling in patient and healthy donor T cells by GUIDE-seq. (**a**) DADA2 patient T cell viability 24-96h after nucleofection with 0-50pmol dsODN/sample for ADA2 locus (measurements performed in quadruplicates). **(b)** DADA2 patient T cell count 24-96h after nucleofection with 0-50pmol dsODN/sample for ADA2 locus (measurements performed in quadruplicates). **(c)** dsODN integration in DADA2 patient T cells with 0-50pmol dsODN/sample for ADA2 locus, assessed by ddPCR (measurements performed in triplicates). **(d)** Schematic representation of the GUIDE-seq experiment. DADA2, APECED and CHH patient and HD PBMCs were thawed and stimulated with IL-2 (120 U/mL), IL-7 (3 ng/μL), IL-15 (3 ng/μL) and soluble CD3/CD28 (15 μL/mL) on day 1, diluted on day 4 and nucleofected on day 5 with 30pmol dsODN and selected RNPs or mock. Cells were cultured in IL-2 (250 U/mL) until sample collection on day 9 for gDNA extraction, ddPCR (dsODN integration) and GUIDE-seq library preparation. GUIDE-seq results in patient and HD T cells for **(e)** ADA2 gRNA #3, **(f)** AIRE gRNA #11, **(g)** RMRP gRNA #9 and **(h)** HEK-site4 gRNA, targeting the endogenous human embryonic kidney HEK site 4. GUIDE-seq results are shown as mismatch plots, where the on-target sequence is depicted at the first line of the table with sequencing read counts on the right. The most abundant off-targets (if applicable) are listed under the target site with their corresponding locations in the genome reported on the left and sequencing read counts on the right. One independent experiment was performed for all sets of data. Bar denotes mean value, error bars represent ± SD. Abbreviations: HD (healthy donor), HDR (homology-directed repair), ddPCR (Droplet Digital PCR), gRNA (guide-RNA), dsODN (double-stranded oligodeoxynucleotide), GUIDE-seq (Genome-wide, Unbiased Identification of DSBs Enabled by Sequencing), RNP (ribonucleoprotein), DADA2 (Deficiency of adenosine deaminase 2), APECED (Autoimmune polyendocrinopathy-candidiasis-ectodermal dystrophy), CHH (Cartilage hair hypoplasia).

### No permanent karyotypic, transcriptomic or proteomic changes in healthy or patient T cells

CRISPR-Cas9 can cause various chromosomal aberrations[46, 47], which increase the risk for malignant transformation and complicate the clinical translation of genome editing. To evaluate the translational potential of our HDR enhancement strategies, we performed a comprehensive search for precancerous lesions in edited T cells.

We first mapped the unintended edits by PacBio long read sequencing from unedited and ADA2 RNP-edited (both groups treated with DMSO or KU0060648) healthy control T cells six days after nucleofection (Supplementary Figure 5a; A-B). We quantified a mean coverage of 25X across the genome, with ∼47% ADA2 HDR editing without enhancer, and ∼67% in combination with KU0060648 (14/31 and 12/18 reads containing the desired edit, respectively; detailed visualization of the cut site available in Supplementary Figure 5a; C). We noted additional on-target indels between ∼3-300bp and a ∼1,2 kb on-target deletion visible in one of the reads of the ADA2-edited, KU0060648 –treated sample. All in all, only one read per edited sample contained no on-target alterations, indicating that almost all cells had been exposed to editing reagents. We found no chromosomal translocations or integrated repair template concatemers on the intended cut site or elsewhere in the genome (Supplementary Figure 5a; C). We also mapped the single nucleotide variants (SNVs), small insertions and deletions outside the cut site. >99% of the detected variation was shared between the experimental conditions, and there were no aberrant mutational signatures[48] (Supplementary Figure 5a; D-E). The findings indicate low risk for malignant transformation; however, combining PacBio with short-read whole genome sequencing could be used to recover low-frequency events.

Single cell RNA sequencing (scRNA-seq) can further evaluate the existence of karyotypic changes and cell states predictive of transformation. To this end, we performed single cell sequencing of full-length mRNA transcripts. For the experiment, DADA2 patient and matched healthy control cells were mock nucleofected or edited with ADA2 RNPs, with both groups treated with KU0060648, IDT-Alt-R or DMSO (total 6 treatment groups, Figure 5A-B). Four days post nucleofection, we sorted single CD4+ and CD8+ cells on plates with 1:1 ratio (Figure 5A). We noted a slight (1.1-1.3X) overrepresentation of CD8+ T cells in cultures after exposure to NHEJ inhibition (Supplementary Figure 5b; A-B). scRNA quantitative real-time PCR (qPCR) detected the presence of the corrected RNA transcript in ∼80% of the RNP-treated and >98% of the NHEJ-inhibited cells, suggesting that all cells had been exposed to editing reagents, and almost all cells harboured at least one corrected allele (Figure 5C, Supplementary Figure 5c; A-B).

**Figure 5.**
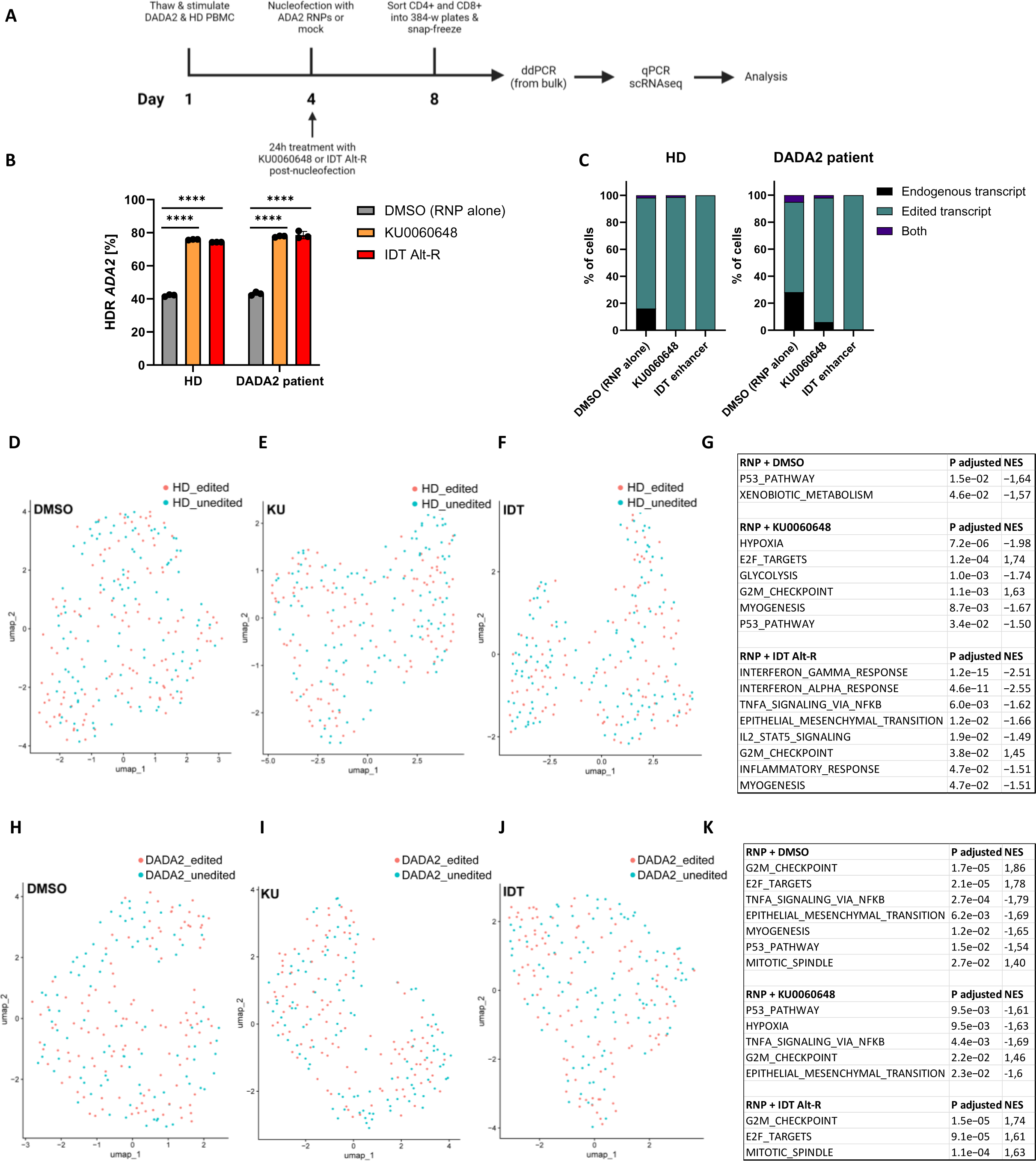
scRNAseq assessment of CRISPR/Cas9 and HDR enhancing compounds in DADA2 patient and HD T cells. (**a**) Outline of the scRNAseq experiment, briefly discussed here: HD and DADA2 patient T cells were thawed and stimulated with IL-2 (120U/mL), IL-7 (3ng/μL), IL-15 (3ng/μL) and soluble CD3/CD28 (15 μL/mL) on day 1 and nucleofected on day 4 with ADA2 RNPs or no cargo. Cells were cultured in IL-2 (250U/mL) and 0.5μM KU0060648, 0.6uM IDT Alt-R enhancer V2 or DMSO for 24h after nucleofection. On day 8, cells were sorted into 384-w plates and gDNA was extracted from the bulk for ddPCR analysis. Sorted cells were processed further for qPCR and scRNA-seq. **(b)** ADA2 HDR editing in HD and DADA2 patient on day 8, assessed by ddPCR (measurements performed in triplicates). **(c)** ADA2 editing outcomes in HD and DADA2 patient, assessed by qPCR of the scRNA-seq libraries with probes to the corrected and uncorrected nucleotide sequence. UMAP plots generated from scRNA-seq for ADA2-edited HD treated with **(d)** DMSO, **(e)** KU0060648 and **(f)** IDT Alt-R enhancer V2, compared to unedited HD (DMSO). **(g)** Hallmark gene set enrichment results for ADA2-edited HD (DMSO, KU0060648 and IDT Alt-R enhancer V2) compared to unedited HD (DMSO). UMAP plots of edited DADA2 patient treated with **(h)** DMSO, **(i)** KU0060648 and **(j)** IDT Alt-R enhancer V2, compared to unedited DADA2 patient (DMSO). **(k)** Hallmark gene set enrichment results for edited DADA2 patient (DMSO, KU0060648 and IDT Alt-R enhancer V2) compared to unedited DADA2 patient. One independent experiment was performed for all sets of data. Bar denotes mean value, error bars represent ± SD. Statistical significance for HDR editing in (b) was assessed by one-way ANOVA with Fisher’s LSD test, where ****p<0.0001. Abbreviations: PBMC (peripheral blood mononuclear cell), HD (healthy donor), IL (interleukin), HDR (homology-directed repair), ddPCR (Droplet Digital PCR), qPCR (quantitative PCR), scRNA-seq (single-cell RNA sequencing), NES (normalized enrichment score), RNP (ribonucleoprotein), UMAP (Uniform Manifold Approximation and Projection), gDNA (genomic DNA).

The analysis of scRNA-seq data recovered no signs of loss of heterozygosity, indicative of lost chromosomal material. The overall transcriptomic effects were minimal, and the edited cells clustered with unedited cells in both healthy control and DADA2 patient (Figure 5D-F, H-J). All edited samples showed a slight downregulation of the p53 response, likely as an adaptive response to the transient p53 upregulation[13] when the ADA2 gene was cut. KU0060648-treated samples displayed additional borderline significant effect on metabolism, likely due to the compound’s bystander effect on PI3K kinase[49]. IDT Alt-R showed downregulation of immune response pathways in the healthy control sample (Figure 5G). In addition, all samples recovered a low frequency non-recurring novel fusion transcripts (Supplementary Table 2). Fusion transcripts that mapped to genes in chromosome 22 were not found in >1 cells per condition. All in all, this data indicates low malignant transformation risk for the corrected DADA2 patient and healthy control T cells, with or without NHEJ inhibition.

To finalize safety profiling, we studied the proteome of the edited and unedited DADA2 and healthy control T cells by mass spectrometry (Supplementary Tables 3-5). We collected the cells seven days after nucleofection to ensure time for altered protein expression (Figure 6A). The only major alteration was the appearance of the ADA2 protein in corrected DADA2 T cells (Figure 6E-G). We found no other statistically significant alterations in samples edited with RNP only, indicating minimal persisting interference (Figure 6E, Supplementary Figure 6C). In samples edited with RNP and NHEJ inhibitors, gene set enrichment analysis[50, 51] identified minor alterations without clear clustering to pathways (Figure 6F-G, Supplementary Figure 6D-E, Supplementary Table 5). None indicated a risk of malignancy for the corrected cells.

**Figure 6.**
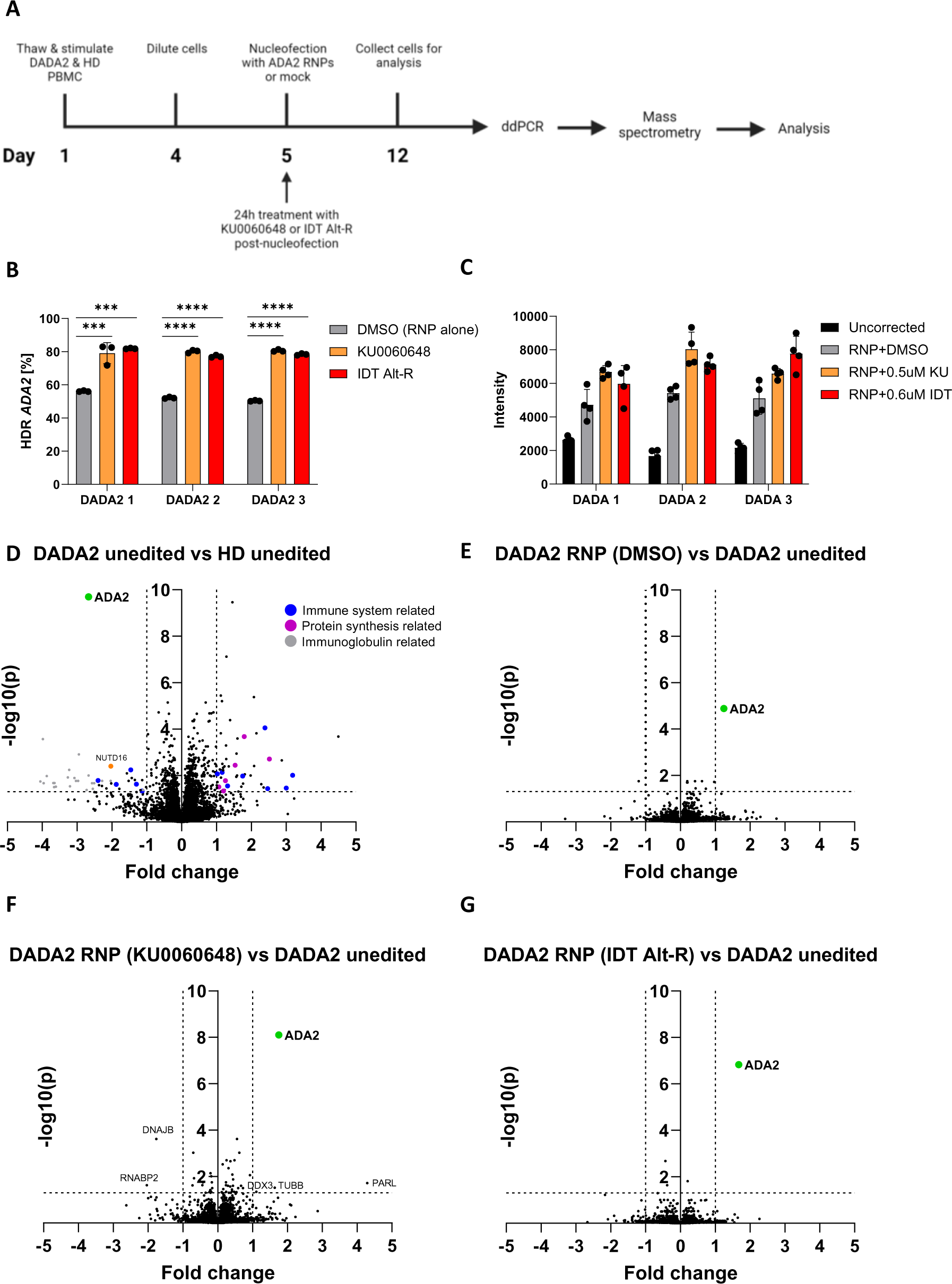
Mass spectrometry analysis of corrected and uncorrected DADA2 patient T cells. (**a**) Outline of the experiment, briefly discussed here: DADA2 patient and HD PBMCs were thawed and stimulated with IL-2 (120 U/mL), IL-7 (3 ng/μL), IL-15 (3 ng/μL) and soluble CD3/CD28 (15 μL/mL) on day 1 and diluted on day 4 for further expansion. Cells were nucleofected with RNPs for ADA2 editing on day 5 and cultured in IL-2 (250 U/mL) and HDR enhancers (0.5 μM KU0060648, 0.6 μM IDT Alt-R enhancer V2) for 24h after nucleofection. Afterwards, cells were cultured in IL-2 (250 U/mL) until they were collected for mass spectrometry and gDNA extraction on day 12. **(b)** Quantification of ADA2 HDR editing in three DADA2 patients treated with HDR enhancers (0.5 μM KU0060648, 0.6 μM IDT Alt-R enhancer V2) or DMSO (RNP alone), assessed by ddPCR (measurements performed in triplicates). **(c)** Abundance of ADA2 protein in DADA2 patients after editing, reported as intensities. **(d)** Comparison of protein expression in unedited (DMSO) DADA2 patients to unedited (DMSO) HDs, assessed by mass spectrometry. (d) Comparison of protein expression in RNP (DMSO) edited DADA2 patients to unedited (DMSO) DADA2 patients, assessed by mass spectrometry. **(e)** Comparison of protein expression in RNP+KU0060648-treated DADA2 patients to unedited (DMSO) DADA2 patients, assessed by mass spectrometry. **(f)** Comparison of RNP+IDT Alt-R enhancer V2 –treated DADA2 patient to unedited (DMSO) DADA2 patients, assessed by mass spectrometry. For (d)-(g), volcano plots were created by reporting fold change of mean protein expression from three DADA2 patients and three HDs on the x axis and –log10 p value on the y axis. One independent experiment was performed for all sets of data. Statistical significance was assessed by one-way ANOVA with Fisher’s LSD test, where ****p<0.0001,***p<0.0002,**p<0.01 and *p<0.05. Bar denotes mean value, error bars represent ± SD. Abbreviations: HD (healthy donor), HDR (homology-directed repair), gDNA (genomic DNA), ddPCR (Droplet Digital PCR), DADA2 (Deficiency of adenosine deaminase 2), RNP (ribonucleoprotein), NUDT (U8 snoRNA-decapping enzyme), DNAJB (DnaJ homolog subfamily B), RNABP2(E3 SUMO-protein ligase RanBP2), DDX3(ATP-dependent RNA helicase),TUBB(Tubulin Beta) and PARL(Presenilin-associated rhomboid-like protein).

### Functional consequences of ADA2 correction

DADA2 is a complex autoinflammatory disease with multiple affected blood cell lineages. The disease hallmark is enhanced INF-γ and TNF-α signalling. Consequently, we saw enhanced TNF-α signaling in unedited DADA2 T cell transcriptomes compared to unedited healthy control (Supplementary Figure 5d; D), suggesting that T cells can, with limitations, be used to model disease pathology. Unedited patient and healthy control T cells clustered separately, and somewhat unexpectedly, patient T cell transcriptomes indicated downregulation of INF-α– and INF-γ responses (Supplementary Figure 5d; A-D). In addition, CD4+ T cells showed up– and CD8+ cells the downregulation of NF-κB and IL-2-STAT5 signaling. Other significant upregulated pathways in patient CD8+ transcriptomes include KRAS signaling and mitotic spindle proteins, likely reflecting enhanced T cell proliferation.

As expected after ADA2 correction, we saw the downregulation of TNF-α signaling in samples corrected with RNP alone, or with RNP in combination with KU0060648. The effects were not visible in cells corrected in the presence of IDT Alt-R, possibly due to the compound interfering with immune signaling pathways (Figure 5H-K). The corrected DADA2 T cell transcriptomes continued to cluster with uncorrected cells. The corrected cells will likely need longer culture and restimulation with appropriate cytokines to show a noticeable shift towards a “healthy” T cell state.

We also compared the expression of >8000 human proteins between three unedited and edited DADA2 patients and healthy controls by mass spectrometry after 12 days in culture (Figure 6C-G). We saw low but detectable ADA2 expression in patients when all MS DIA runs were searched together; however, when searched alone no ADA2 was detected, suggesting very low or no ADA2 expression in the patients (Figure 6C, Supplementary Table 4). In addition, the DADA2 T cells showed downregulation of several proteins implicated in inflammatory response, as well as lowered expression of the mRNA decapping enzyme NUTD16 (Figure 6D, Supplementary Tables 4 and 5). Consequently, the proteins of the translational machinery were upregulated, along with several adaptive immune response proteins. We also detected cytoplasmic immunoglobulins, which we attribute to residual B cells in the samples, as we saw no immunoglobulin transcripts in the scRNA-seq data where T cells were pre-sorted using flow cytometry. Upon DADA2 correction, we saw up to two-fold increase in ADA2 expression. In controls, ADA2 expression decreased upon editing in two of the three controls, either due to on-target NHEJ deletions or due to addition of silent SNVs (Supplementary Figure 6B).

### Gene correction rescues T cell proliferation in Cartilage-Hair Hypoplasia

Mutations in RMRP cause Cartilage-Hair hypoplasia (CHH), a syndromic immunodeficiency with defective T cell proliferation[52]. We thus evaluated the patient T cell proliferative capacity in response to mutation correction. We further speculated that the corrected patient T cells would outgrow their uncorrected counterparts, and consequently the frequency of corrected alleles would increase in DNA samples taken during prolonged CHH T cell culture.

To test this, we first corrected RMRP in T cells from three CHH patients and measured HDR correction levels at 4-, 7-, and 14 days post-nucleofection by amplicon sequencing (Figure 7A). We noted up to 50% correction at day 4, with increases up to 70% at 14 days post-nucleofection, with individual variation. Consequently, we chose to assess T cell proliferative capacity 14 days post-nucleofection and enhance RMRP correction by treating cells with KU0060648 for the first 24 hours after nucleofection, as we observed no increased toxicity from NHEJ inhibition (Supplementary Figure 3K). We performed CFSE based T cell proliferation assay in four corrected and uncorrected CHH patients (Figure 7B) 14 days after nucleofection (day 20 in cell culture). We saw improved CD4+ T cell proliferation in all corrected patients (Figure 7C) and improved CD8+ T cell proliferation in all but one patient (Figure 7D). In conclusion, genomic correction of the RMRP enhances T cell proliferation, which leads to selective growth advantage for the corrected cells.

**Figure 7.**
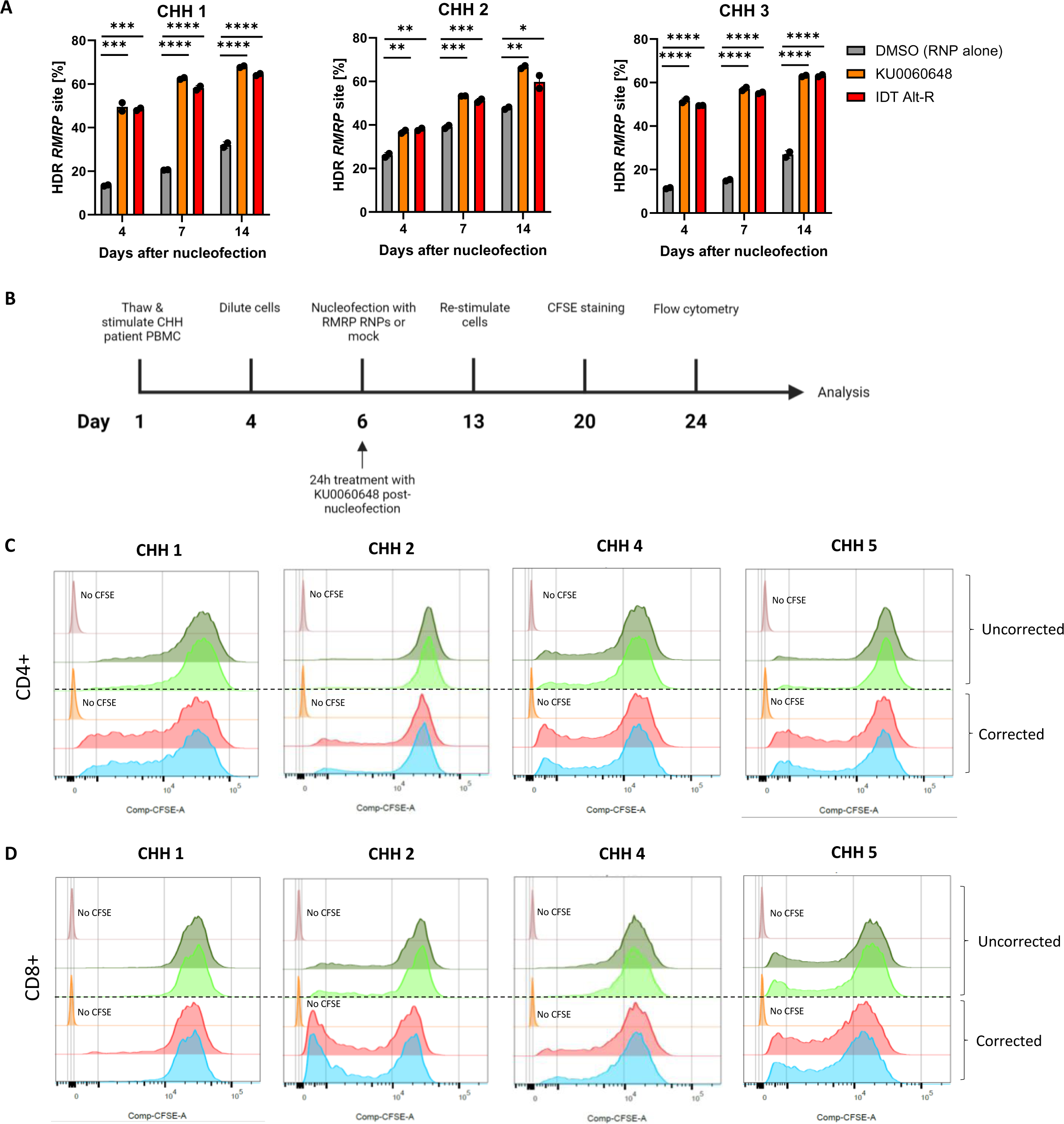
T cell proliferation assay in Cartilage-hair hypoplasia patients. (**a**) Quantification of RMRP HDR editing levels in three corrected CHH patients at 4, 7 and 14 days after nucleofection with concentration-optimized HDR enhancing small molecules (0.5 μM KU0060648, 0.6 μM IDT Alt-R enhancer V2) or DMSO (RNP alone). HDR levels were assessed by amplicon sequencing (measurements performed in duplicates). **(b)** Outline of the experiment, briefly discussed here: CHH patient PBMCs were thawed and stimulated with IL-2 (120 U/mL), IL-7 (3 ng/μL), IL-15 (3 ng/μL) and soluble CD3/CD28 (15 μL/mL) on day 1 and diluted on day 4 for further expansion. Cells were nucleofected with RNPs for RMRP correction or mock on day 6 and cultured in IL-2 (250 U/mL) and 0.5 μM KU0060648 for 24h after nucleofection. Afterwards, cells were cultured in IL-2 (250 U/mL) until they were re-stimulated on day 13 with the same setup as on day 1. Cells were stained with CFSE on day 20 and cultured in IL-2 (250 U/mL) for four days. On day 24, cells were stained for flow cytometry to assess T cell proliferation. **(c)** CD4+ T cell proliferation assessed by flow cytometry in corrected and uncorrected CHH patients after CFSE staining. **(d)** CD8+ T cell proliferation assessed by flow cytometry in corrected and uncorrected CHH patients after CFSE staining. One independent experiment was performed for all sets of data. The patient number corresponds to the numbering in the supplementary patient table. Bar denotes mean value, error bars represent ± SD. Statistical significance was assessed by one-way ANOVA with Fisher’s LSD test, where ****p<0.0001,***p<0.0002,**p<0.01 and *p<0.05. Abbreviations: CHH (Cartilage-hair hypoplasia), HD (healthy donor), HDR (homology-directed repair), gDNA (genomic DNA), CFSE (Carboxyfluorescein succinimidyl ester), RNP (ribonucleoprotein).

## DISCUSSION

In this study, we evaluated and combined several novel and previously published methods to develop a pipeline for enhanced HDR based gene correction, targeting Inborn Errors of Immunity. We demonstrate up to 80% biallelic correction in T cells, with individual variation[13, 32, 35, 36], and functional correction of the model disease phenotypes. Since the pipeline is suitable for correcting SNV and small indels multiple genes and the protocol reagents can be upgraded into a GMP compatible format, it can be further developed into a platform technology for generating corrected T cells. Corrected autologous T cell transplants could further be developed to a salvage therapy for IEI patients with isolated T cell defects, who are not suitable for stem cell transplants[11].

The primary concern with therapeutic gene editing technology is safety. We could not detect precancerous genomic lesions when the model conditions were evaluated with single cell RNA sequencing, PacBio long-read whole genome sequencing, GUIDE-Seq and mass spectrometry proteomic profiling. This is in line with other reports validating the safety of human primary cell editing[53]. CRISPR-Cas9 cutting can result in a variety of persistent on-target structural chromosomal changes[47, 54–57] in 5-20% frequency, with optimized culture conditions decreasing the events[47]. We detected no chromosomal aberrations, possibly because the cells with larger abnormalities permanently arrested and disappeared below detection limit at the assay timepoint[47, 57]. The addition of DNA-PKcs inhibition by KU0060648 did not lead to persistent changes in cell state or major genomic alterations. When preparing this manuscript, a third DNA-PKcs inhibitor (AZD7648) was identified[58,59] independent of this study, strengthening our observation of DNA-PKcs inhibition as an effective and safe strategy for HDR enhancement in T cells and hematopoietic stem cells.

Our correction strategy introduces 2-4 silent SNVs along with the correction of the pathogenic variant. The strategy prevents CRISPR re-cutting after successful HDR repair and thus improves precise correction levels[29, 38]. Additionally, it allows very accurate and rapid editing quantification by droplet digital PCR. However, the introduction of extra variation within the coding sequence may interfere with the expression of the corrected gene or affect the regulation of more distant genes. To ensure minimal interference with gene function and regulation, we advise prioritizing SNVs that are part of normal human variation and located in evolutionarily less conserved regions.

Inborn Errors of Immunity currently encompass mutations in >500 different genes, with novel disease genes discovered monthly. To offer gene therapy for most IEI patients, a combination of correction strategies – therapeutic cDNA-knock in for more common conditions, and the combinations of base-prime– and HDR-based correction for ultra-rare diseases – is necessary. In addition, the possibility for diverse correction strategies is important because many IEI genes affect DNA repair and cell danger signalling pathways, which can decrease editing efficiency and increase risk for unintended genomic events, regardless of the editing strategy. In future studies, the efficacy and safety of our suggested correction strategy should be studied diverse IEI patient samples along with healthy controls to understand how pathology and normal human variation affect genome editing outcomes.

In conclusion, we present a non-viral T cell SNV correction protocol that has the potential to be scaled up to a platform technology to correct diverse pathogenic single nucleotide variants, small deletions, and insertions in Inborn Errors of Immunity. Our pipeline is compatible for GMP scale-up and is also useful for basic studies of normal and pathological variation of the human immune system.

## AUTHOR CONTRIBUTIONS

**K.M.** performed most of the experiments and wrote the manuscript. **S.Ko.** & **ME** performed scRNA-seq experiments. **Z.L.** designed CRISPR reagents and performed GUIDE-seq optimization. **K.L.**, **E.T.** & **E.V.** performed bioinformatic & data analysis. **A.K.** performed GUIDE-seq and PacBio experiments. **S.Ke., A.T & M.V** performed mass spectrometry experiments and data analysis. **G.R.** designed CRISPR reagents, amplicon sequencing panel and performed experiments. **F.H.H.** performed gRNA screening and BG-coupled ssODN experiments. **B.O.L.** performed CRISPR optimization experiments in T cells. **T.J.G** performed flow cytometry experiments. **C.W.E.** & **P.K.** performed gRNA screening in CD34+ HSPCs. **N.F. & M.S.** performed library preparation for amplicon sequencing **T.M.M.** obtained cord blood for HSPC isolation. **J.S. & J.O.** supervised experiments. **V.G., E.L., C.S.J., T.H. & T.M** provided clinical care for the patients & obtained samples. **S.D.K.** designed scRNA-seq, flow cytometry and cell sorting experiments, performed & supervised research and wrote the manuscript. **E.H.** supervised the study and wrote the manuscript. All authors read and approved the manuscript.

## Supporting information

Supplementary Table 1_Patient table

Supplementary Table 2_Fusion Detection

Supplementary Table 3_Mass spectrometry data_vs HD_unedited

Supplementary Table 4_Mass spectrometry data_vs MUT_unedited

Supplementary Table 5_ Mass spectrometry significant hits DADA2 patients

Supplementary Table 6_Barcodes_for_Single_cell_sequencing

## ACKNOWLEDGEMENTS

We thank all patients and families for their participation in the study. We thank Karolinska Institute Protein Science Facility for manufacturing the Cas9 protein. Research Council of Norway, Norway Health South-East Region, Norwegian Cancer Society, and the Swedish Childhood Cancer Fund (Barncancerfonden) funded the study.

---

**Supplementary Figure 1.**
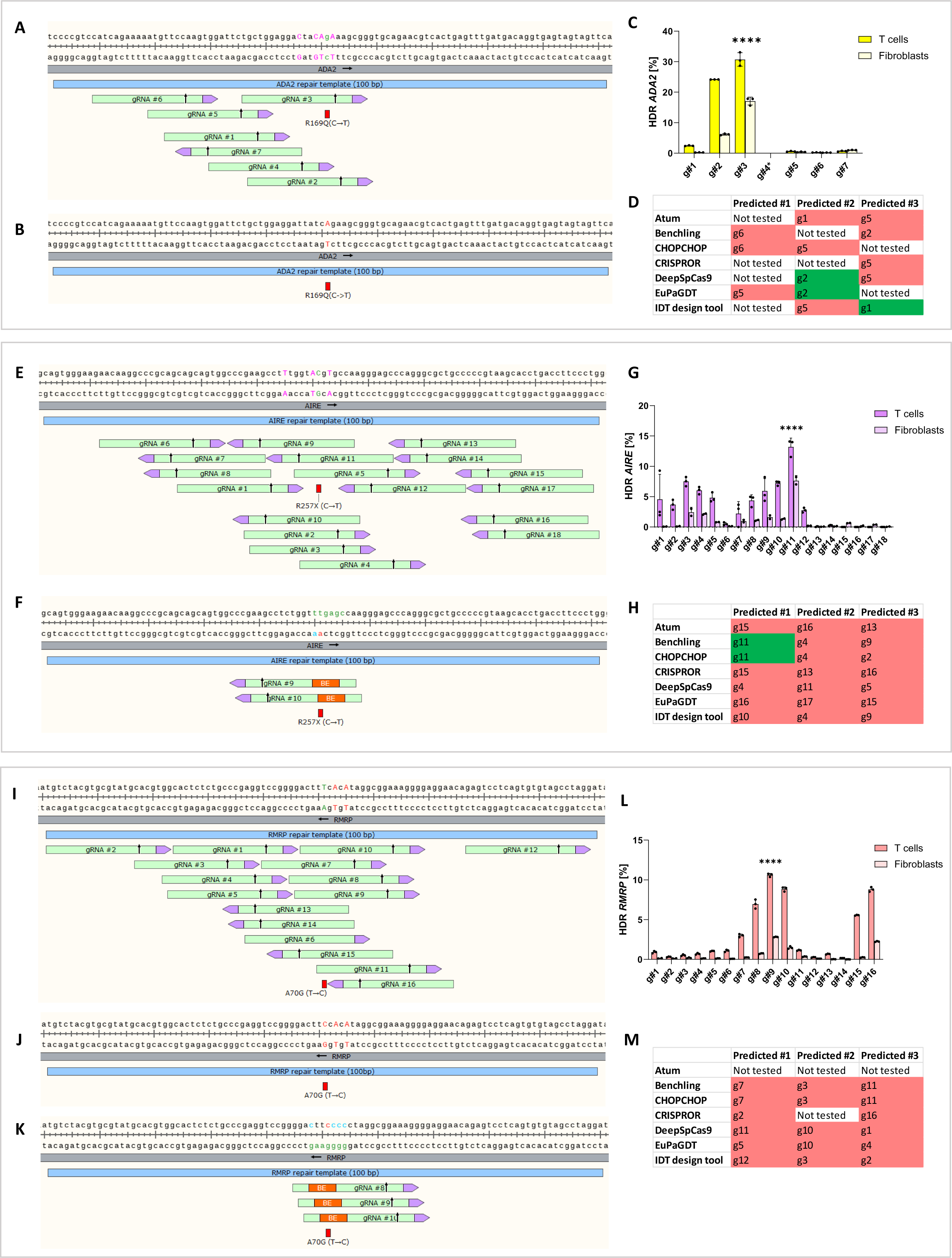
Repair template design and guide-RNA screening for ADA2, AIRE and RMRP. (**a**) Schematic representation ADA2 gRNA design and SNP strategy. Correction of pathogenic mutation (red) is shown in green uppercase letter and silent SNPs as pink uppercase letters. **(b)** Schematic representation of ADA2 mutant site, marked with red uppercase letter, with no possible base editing gRNAs. **(c)** ADA2 gRNA screening in patient T cells and fibroblasts, assessed by amplicon sequencing (measurements performed in triplicates). **(d)** Comparison of three best gRNAs identified by *in silico* gRNA design tools to *in vitro* validated gRNA screening results from DADA2 patient T cells, where accurate predictions are shown in green, incorrect predictions in red and gRNAs that were designed by the tools but not assessed *in vitro* in white. **(e)** Schematic representation AIRE gRNA design and SNP strategy, as explained for (a). **(f)** Schematic representation of AIRE mutant site, where the edited pathogenic mutation nucleotide is marked with red, showing possible A→G base editing gRNAs. The nucleotide positions which fall within the editing window span of BE guides are shown in green and the bystander edits are shown in blue. **(g)** AIRE gRNA screening in patient T cells and fibroblasts, assessed by amplicon sequencing (measurements performed in triplicates for T cells and duplicates for fibroblasts). **(h)** Comparison of three best gRNAs identified by *in silico* gRNA design tools to *in vitro* validated gRNA screening results from APECED patient T cells, where accurate predictions are shown in green, incorrect predictions in red. **(i)** Schematic representation RMRP gRNA design and SNP strategy, as explained for (a). As RMRP is noncoding, non-silent SNPs, marked in red uppercase letters, were added in the repair strategy for early experiments. **(j)** Schematic representation of RMRP SNP strategy for editing WT cells. As RMRP is noncoding, non-silent SNPs, marked in red uppercase letters, were added in the repair strategy for early experiments. **(k)** Schematic representation of RMRP mutant site, where the edited pathogenic mutation nucleotide is marked with red, showing possible A→G base editing gRNAs. The nucleotide positions which fall within the editing window span of BE guides are shown in green and the bystander edits are shown in blue. **(l)** RMRP gRNA screening in patient T cells and fibroblasts, assessed by amplicon sequencing (measurements performed in triplicates). **(m)** Comparison of three best gRNAs identified by *in silico* gRNA design tools to *in vitro* validated gRNA screening results from CHH patient T cells, where accurate predictions are shown in green, incorrect predictions in red and gRNAs that were designed by the tools but not assessed *in vitro* in white. One independent experiment was performed for all sets of data. Statistical significance of highest HDR for a given gRNA was assessed by one-way ANOVA with Fisher’s LSD test, where ****p<0.0001. Bar denotes mean value, error bars represent ± SD. Abbreviations: HDR (homology-directed repair), gRNA (guide-RNA), HD (healthy donor), SNP (single nucleotide polymorphism), nt (nucleotide), DADA2 (Deficiency of adenosine deaminase 2), APECED (Autoimmune polyendocrinopathy-candidiasis-ectodermal dystrophy), CHH (Cartilage hair hypoplasia).

**Supplementary Figure 2.**
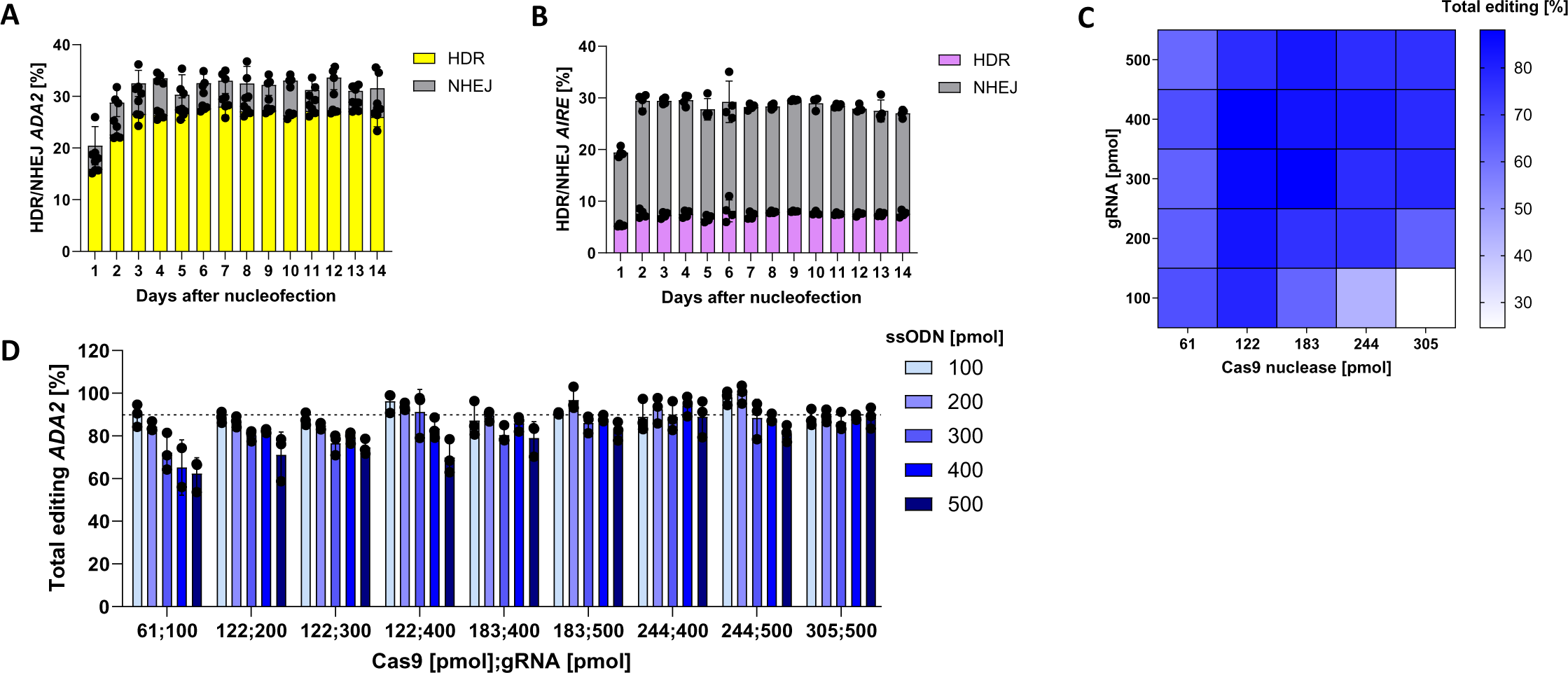
Optimization of CRISPR reagents and nucleofection in healthy donor T cells. (**a**) ADA2 HDR and NHEJ editing in HD T cells 1-14 days after nucleofection, measured by ddPCR (measurements performed in triplicates). **(b)** AIRE HDR and NHEJ editing in HD T cells 1-14 days after nucleofection, measured by ddPCR (measurements performed in triplicates). **(c)** ADA2 total editing (reported as the sum of HDR and NHEJ) in HD T cells with Cas9 nuclease at 61-305pmol, gRNA at 100-500pmol and ssODN at 100pmol/sample, measured by ddPCR (measurements performed in triplicates). **(d)** ADA2 total editing (reported as the sum of HDR and NHEJ) in HD T cells with selected combinations of RNPs with ssODN at 100-500pmol/sample, measured by ddPCR (measurements performed in triplicates). Dashed line indicates mean measurements for Cas9 nuclease at 61pmol, gRNA at 100pmol and ssODN at 100pmol/sample. One independent experiment was performed for all sets of data except for (c)-(d) where one out of three representative experiments is shown. Bar denotes mean value, error bars represent ± SD. Abbreviations: HD (healthy donor), gRNA (guide-RNA), ssODN (single-stranded oligodinucleotide), NHEJ (non-homologous end joining), HDR (homology-directed repair), ddPCR (Droplet Digital PCR), RNP (ribonucleoprotein).

**Supplementary Figure 3.**
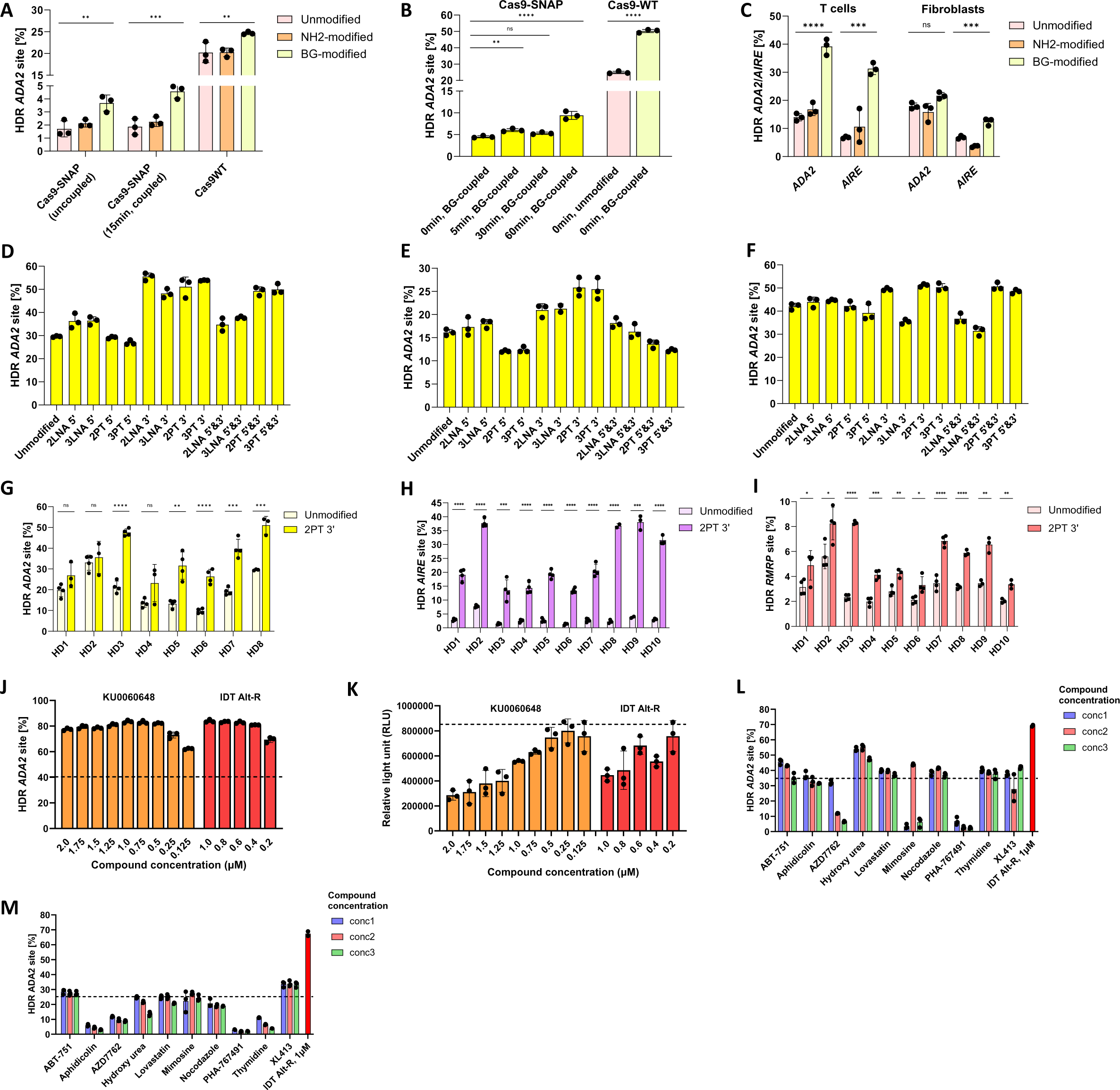
Assessing HDR improvement strategies in healthy donor primary cells. (**a**) Quantification of ADA2 HDR editing in HD fibroblasts with unmodified, NH2-or BG-modified ssODNs tested with Cas9-SNAP (uncoupled or coupled) or Cas9WT nuclease (uncoupled), measured by ddPCR (measurements performed in triplicates). **(b)** Quantification of ADA2 HDR editing in HD T cells with unmodified (pink bar) or BG-modified (bright and pale-yellow bars) ssODNs with Cas9-SNAP or Cas9WT nuclease, measured by ddPCR (measurements performed in triplicates). **(c)** Quantification of ADA2 and AIRE HDR editing in HD T cells and fibroblasts, respectively, with unmodified, NH2-or BG-modified ssODNs with Cas9WT nuclease, measured by ddPCR (measurements performed in triplicates). Quantification of ADA2 HDR editing with LNA– and PT-modified ssODNs in **(d)** HD T cells **(e)**, HD fibroblasts and **(f)** HD CD34+ HSPCs, measured by by ddPCR (measurements performed in triplicates). Quantification of HDR editing in 8-10 healthy T cell donors with position-optimized ssODNs with unmodified and 2PT 3’ modified ssODNs for **(g)** ADA2, **(h)** AIRE and **(i)** RMRP, measured by ddPCR (measurements performed in triplicates and quadruplicates depending on the donor). Effect of HDR enhancing compounds at selected concentrations (0.125-2 μM KU0060648, 0.2-1 μM IDT Alt-R enhancer V2) in HD T cells on **(j)** ADA2 HDR editing, measured by ddPCR (measurements performed in triplicates), where dashed line indicates the mean value for DMSO (RNP alone), and **(k)** cell viability 96h after nucleofection, measured by CellTiter-Glo (measurements performed in triplicates). Quantification of ADA2 HDR editing in HD T cells with cell cycle inhibitors at three concentrations (conc1-conc3) applied **(l)** 24h pre– and **(m)** 24h post nucleofection, measured by ddPCR (measurements performed in triplicates). Dashed line indicates the mean value for DMSO (RNP alone). A single experiment was performed for all sets of data except for (j) and (k) where three independent experiments were performed, and the representative experiment is shown. Statistical significance was assessed by one-way ANOVA with Fisher’s LSD test, where ****p<0.0001, ***p<0.0002, **p<0.001 and *p<0.01. Bar denotes mean value, error bars represent ± SD. Abbreviations: HD (healthy donor), ssODN (single-stranded oligodinucleotide), HDR (homology-directed repair), ddPCR (Droplet Digital PCR), RNP (ribonucleoprotein), LNA (locked nucleic acid), PT (phosphorothioate), BG (benzylguanine).

**Supplementary Figure 4.**
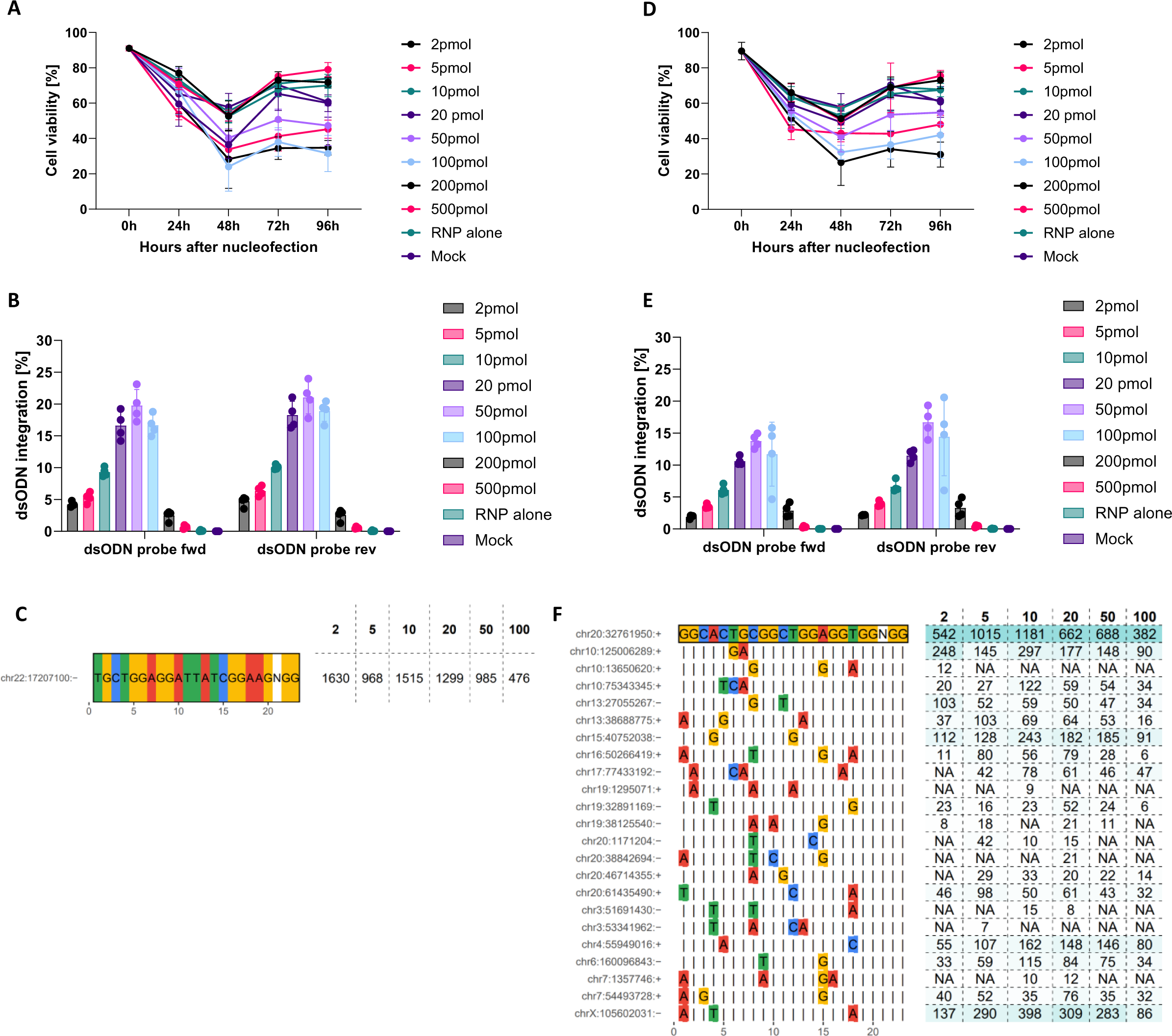
GUIDE-seq optimization in healthy donor T cells. (**a**) HD T cell viability 24-96h after nucleofection with 0-500pmol dsODN/sample for ADA2 locus (measurements performed in quadruplicates). **(b)** dsODN integration in HD T cells with 0-500pmol dsODN/sample for ADA2 locus, assessed by ddPCR (measurements performed in quadruplicates). **(c)** GUIDE-seq mismatch plots for ADA2 gRNA #3 in HD T cells with dsODN at 2-100pmol/sample. On-target sequence is reported at the top of the table and sequencing reads for each dsODN concentration at the right. **(d)** HD T cell viability 24-96h after nucleofection with 0-500pmol dsODN/sample for HEK-site4 locus (measurements performed in quadruplicates). **(e)** dsODN integration in HD T cells with 0-500pmol dsODN/sample for HEK-site4 locus, assessed by ddPCR (measurements performed in quadruplicates). **(f)** GUIDE-seq mismatch plot for HEK-site4 gRNA in HD T cells with dsODN at 2-100pmol/sample. The most abundant off-targets are listed under the target site with their corresponding locations in the genome reported on the left and sequencing read counts on the right. Coloured bases of off-targets indicate mismatches with the on-target site One independent experiment was performed for all sets of data. Bar denotes mean value, error bars represent ± SD. Abbreviations: HD (healthy donor), HDR (homology-directed repair), ddPCR (Droplet Digital PCR), gRNA (guide-RNA), dsODN (double-stranded oligodeoxynucleotide), GUIDE-seq (Genome-wide, Unbiased Identification of DSBs Enabled by Sequencing), RNP (ribonucleoprotein).

**Supplementary Figure 5a.**
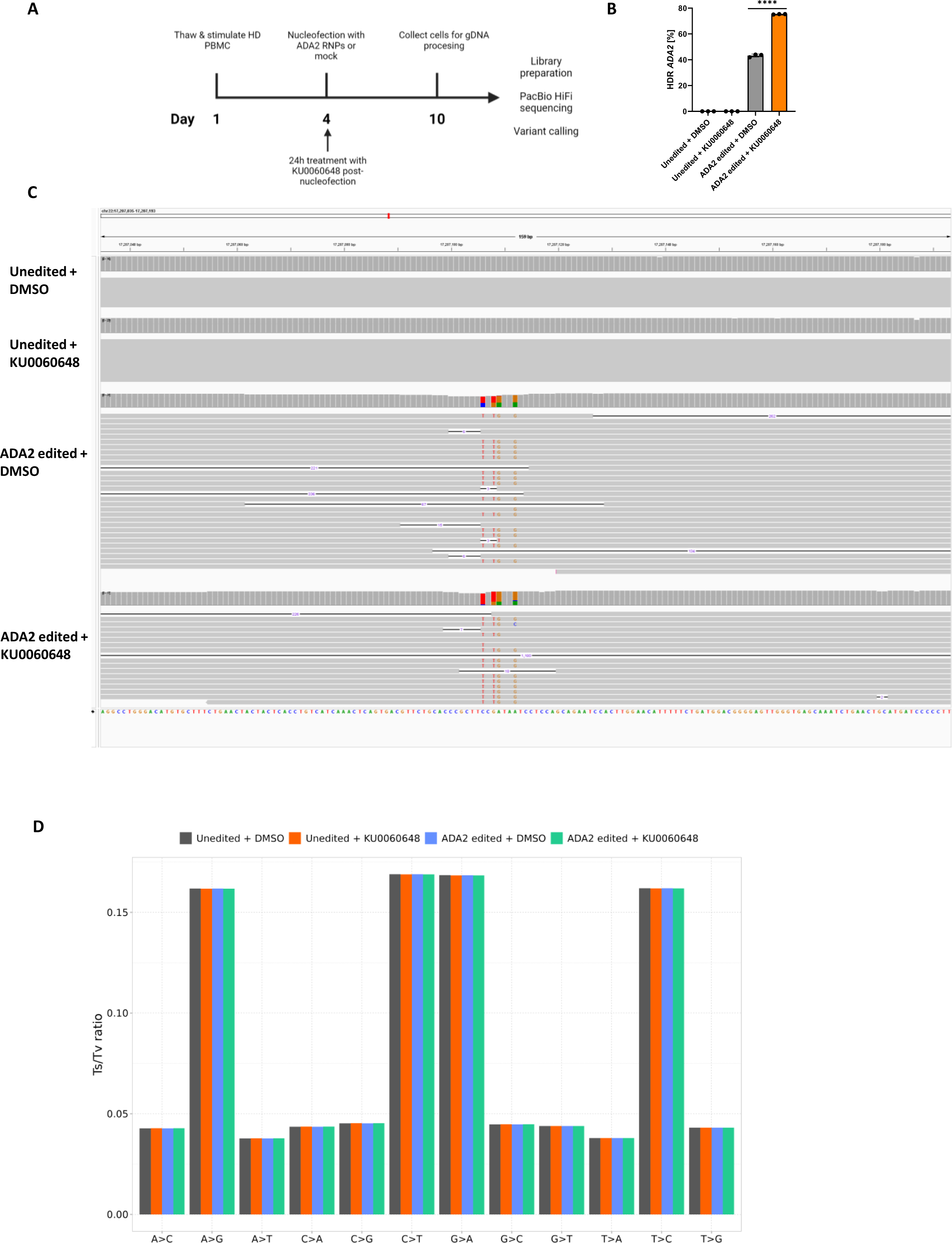
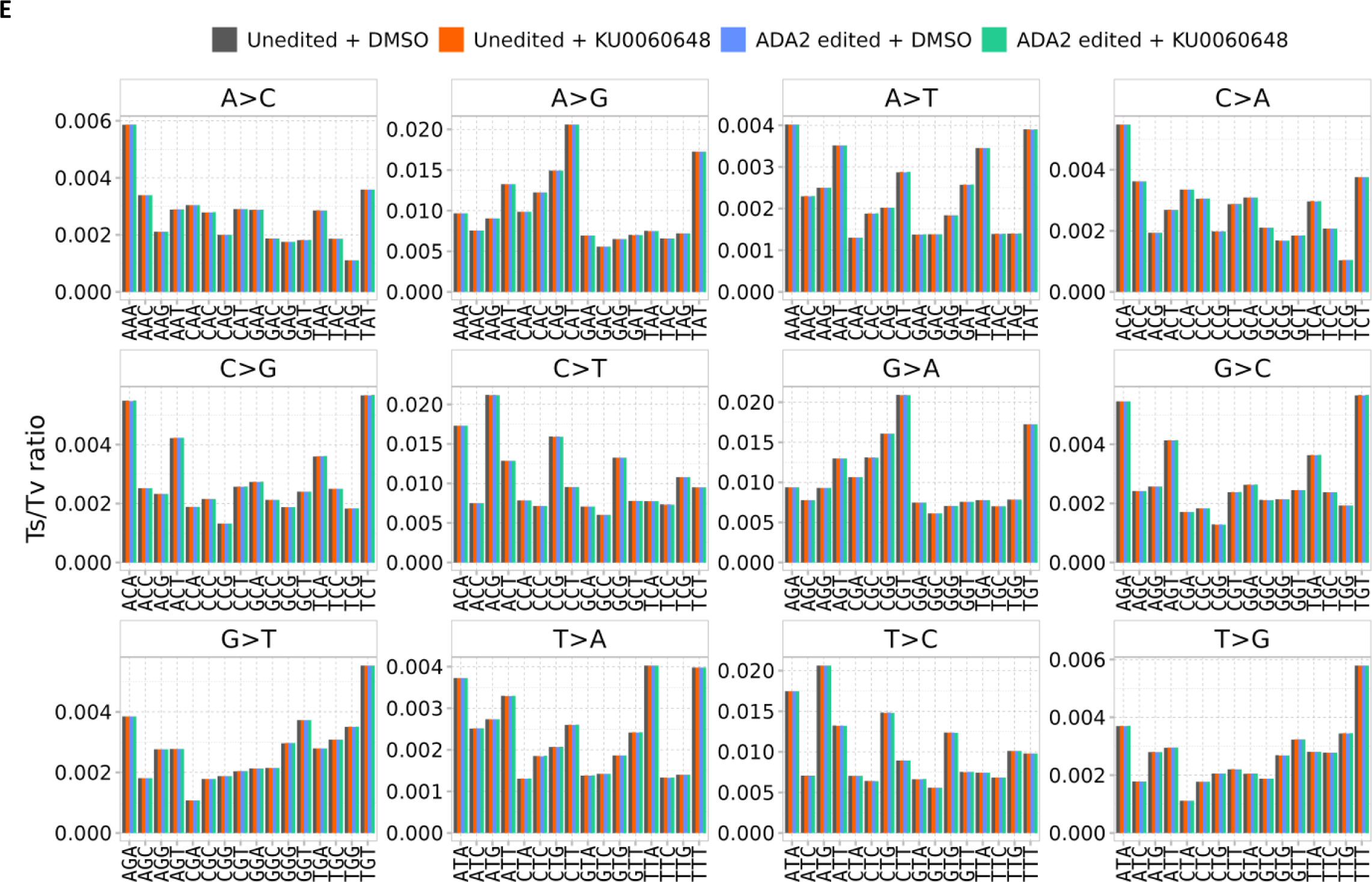
Whole genome sequencing of unedited and ADA2 edited HD T cells. (**a**) Outline of the WGS experiment, briefly discussed here: HD T cells were thawed and stimulated with IL-2 (120 U/mL), IL-7 (3 ng/μL), IL-15 (3 ng/μL) and soluble CD3/CD28 (15 μL/mL) on day 1 and nucleofected on day 4 with ADA2 RNPs or mock. Cells were cultured in IL-2 (250 U/mL) and 0.5 μM KU0060648 or DMSO for 24h after nucleofection and collected on day 10 of the pipeline. gDNA from samples was processed for ddPCR and PacBio sample preparation, followed by PacBio HiFi sequencing and analysis. **(b)** ADA2 HDR editing levels, assessed by ddPCR (measurements performed in triplicates. **(c)** IGV view of HiFi PacBio reads on the on-target ADA2 site. HDR reads contain four SNVs at the same time: C>T, G>T, A>G and A>G. No editing is present in the unedited samples. **(d)** Transition transversion ratio plot showing no difference between edited and unedited samples, showing no global CRISPR toxicity. **(e)** Mutational signature by codon shows no differences between edited and unedited samples. One independent experiment was performed for all sets of data. Bar denotes mean value, error bars represent ± SD. Statistical significance was assessed by one-way ANOVA with Fisher’s LSD test, where ****p<0.0001. Abbreviations: WGS (whole genome sequencing), PBMC (peripheral blood mononuclear cell), HD (healthy donor), IL (interleukin), gRNA (guide-RNA), ssODN (single-stranded oligodinucleotide), HDR (homology-directed repair), ddPCR (Droplet Digital PCR), RNP (ribonucleoprotein).

**Supplementary Figure 5b.**
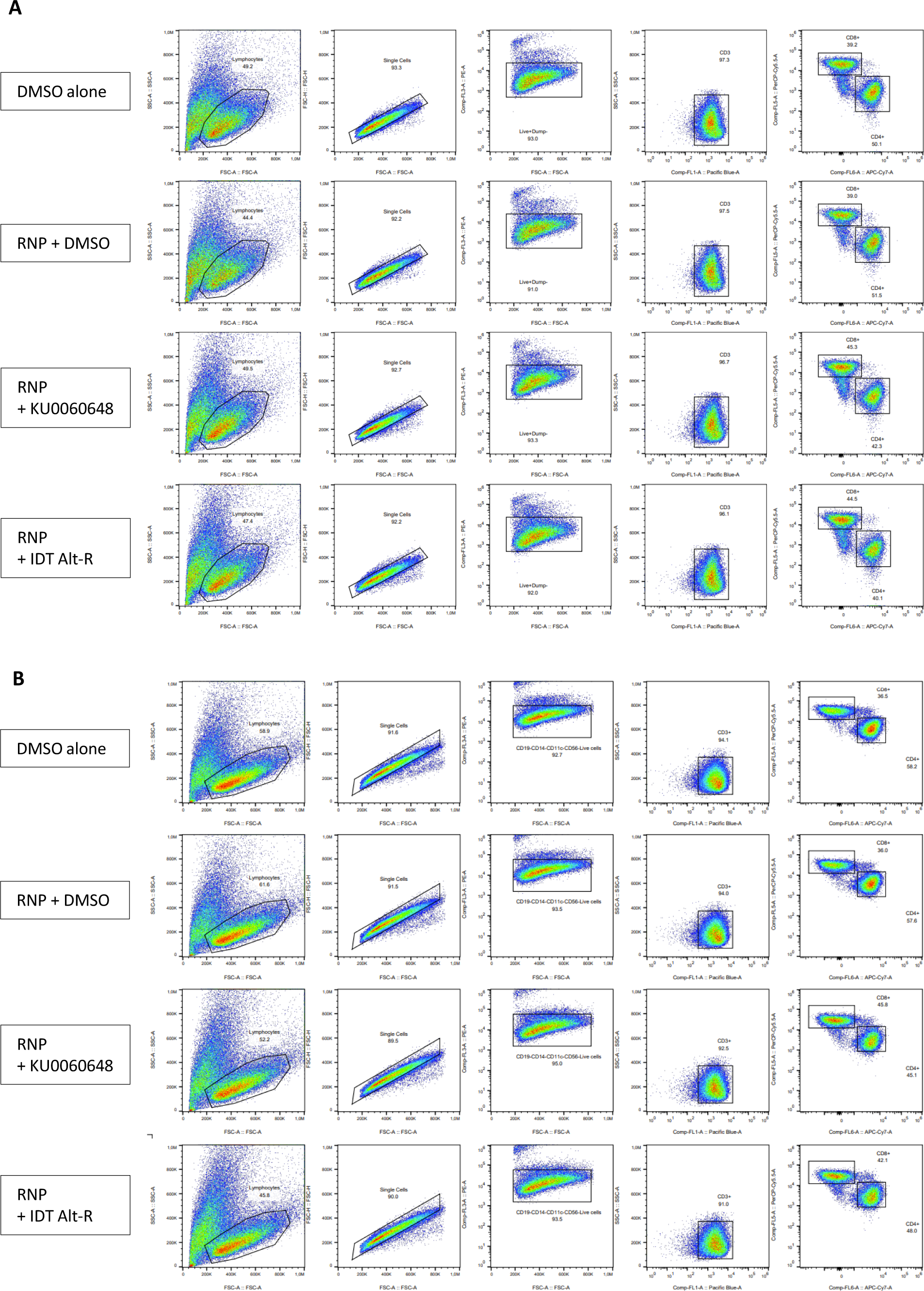
Panel setup for fluorescence-activated cell sorting of CD4+ and CD8+ T cells in HD and DADA2 patient. FACS gating strategy of unedited and ADA2 edited CD4+ and CD8+ T cells in **(a)** HD and **(b)** DADA2 patient. Cells were nucleofected on day 4 of the pipeline and treated with 0.5uM KU0060648, 0.6uM IDT Alt-R enhancer V2 or DMSO for 24h after nucleofection. Samples were collected for FACS on day 8 of the pipeline. One independent experiment was performed for all experiments except (b), where one out of three representative experiments is shown. Abbreviations: HD (healthy donor), DADA2 (Deficiency of adenosine deaminase 2), FACS (fluorescence-activated cell sorting), RNP (ribonucleoprotein).

**Supplementary Figure 5c.**
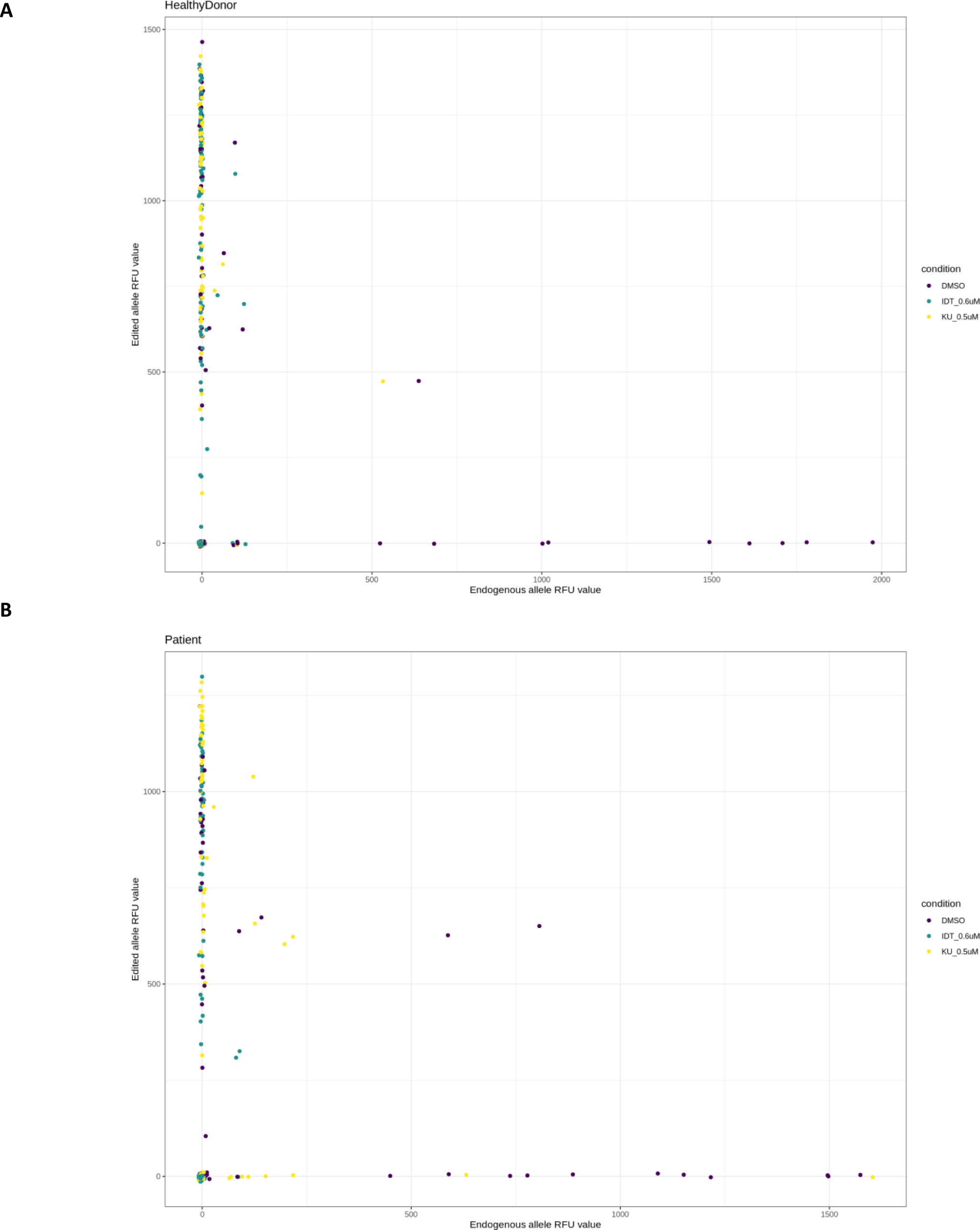
qPCR plots of edited DADA2 patient and healthy donor. (**a**) qPCR plots for edited HD and (b) DADA2 patient. Plots show RFU values of edited allele on y axis and endogenous allele on x axis. Each dot is measurement from a single cell. Dots are colored by which condition the cells underwent editing, DMSO in purple, 0.6uM IDT in green and 0.5uM KU in yellow. Abbreviations: qPCR (quantitative PCR), DADA2 (Deficiency of adenosine deaminase 2).

**Supplementary Figure 5d.**
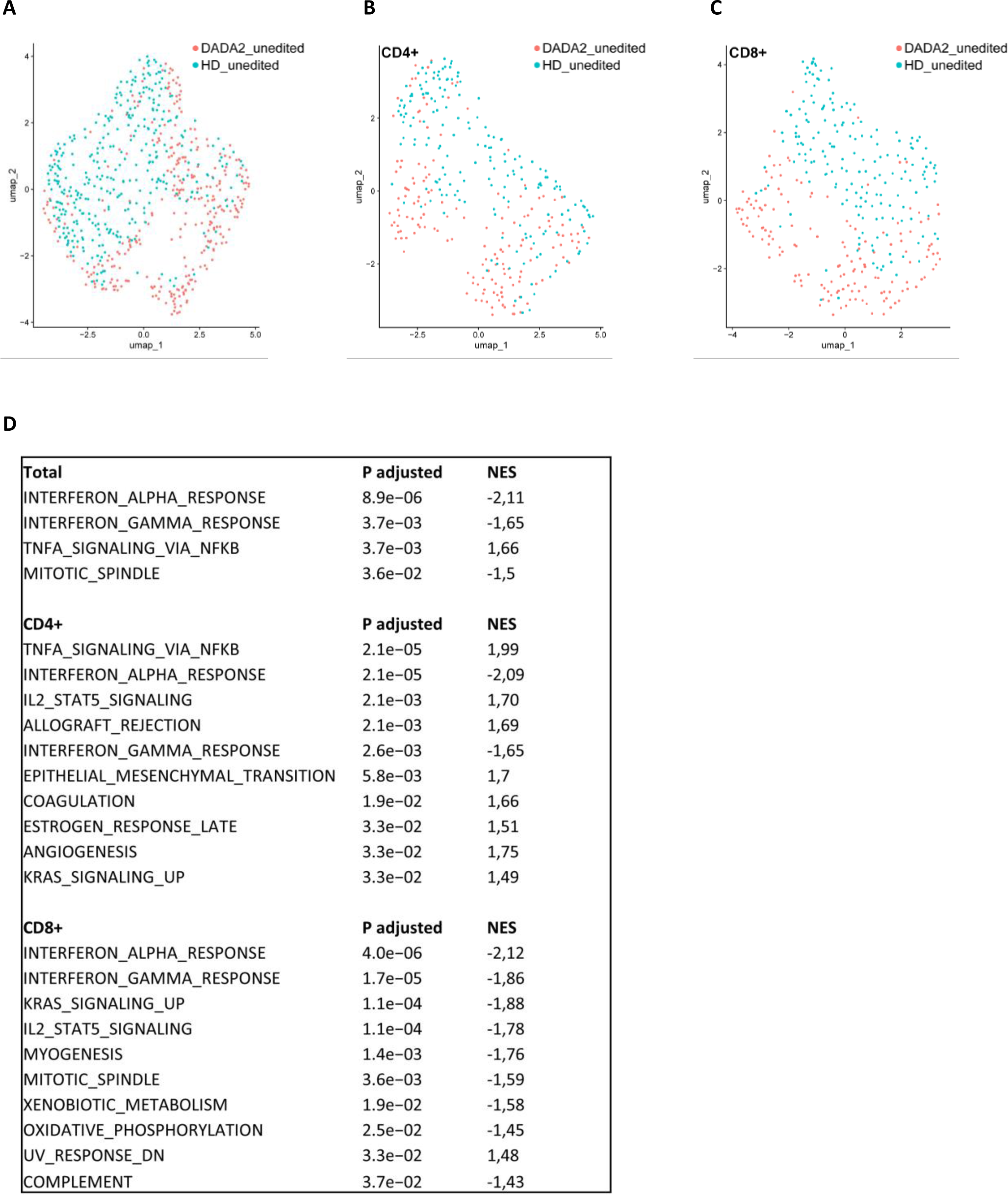
scRNAseq assessment of total, CD4+ and CD8+ T cells in DADA2 patient and HD T cells. UMAP plots generated from scRNA-seq of unedited DADA2 patient for **(a)** total, **(b)** CD4+ and **(c)** CD8+ T cells, compared to unedited HD (DMSO). **(d)** Hallmark gene set enrichment results for unedited DADA2 patient (total, CD4+ and CD8+) compared to unedited HD (DMSO). Abbreviations: HD (healthy donor), scRNA-seq (single-cell RNA sequencing), NES (normalized enrichment score), UMAP (Uniform Manifold Approximation and Projection).

**Supplementary Figure 6.**
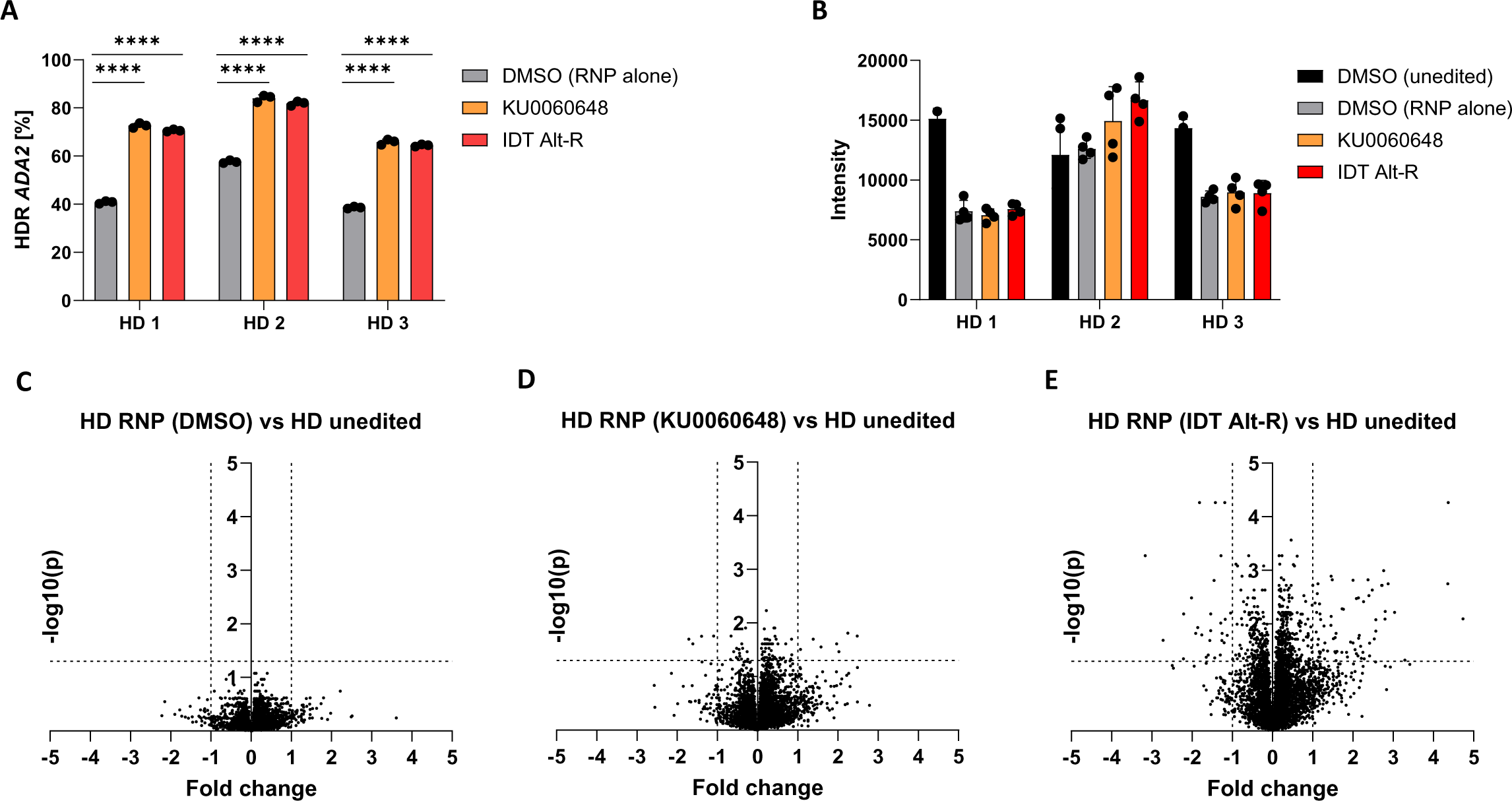
Mass spectrometry analysis of ADA2 edited and unedited healthy donor T cells. (**a**) Quantification of ADA2 HDR editing in three healthy donors with selected HDR enhancers (0.5μM KU0060648, 0.6μM IDT Alt-R enhancer V2) or DMSO (RNP alone), assessed by ddPCR (measurements performed in triplicates). **(b)** Abundance of ADA2 protein in healthy donors after mock and ADA2 RNP editing, reported as mass spectrometry intensities. **(c)** Comparison of protein expression in RNP (DMSO) edited healthy donors to unedited (DMSO) healthy donors, assessed by mass spectrometry. **(d)** Comparison of protein expression in RNP+KU0060648-treated healthy donors to unedited (DMSO) healthy donors, assessed by mass spectrometry. **(e)** Comparison of RNP+IDT Alt-R enhancer V2 –treated healthy donors to unedited (DMSO) healthy donors, assessed by mass spectrometry. For (c)-(e), volcano plots were created by reporting fold change of mean protein expression from three healthy donors on the x axis and –log10 p value on the y axis. One independent experiment was performed for all sets of data. Statistical significance was assessed by one-way ANOVA with Fisher’s LSD test, where ****p<0.0001. Bar denotes mean value, error bars represent ± SD. Bar denotes mean value, error bars represent ± SD. Abbreviations: ADA2 (adenosine deaminase 2), (HD (healthy donor), HDR (homology-directed repair), ddPCR (Droplet Digital PCR), RNP (ribonucleoprotein), NUDT.

