## Supplemental Methods for "T cell correction pipeline for Inborn Errors of Immunity"

**SUPPLEMENTARY MATERIALS AND METHODS**

**Patient and healthy donor sample collection**

We obtained peripheral blood, cord blood, and fibroblasts from human donors. The study was conducted per the principles of the Helsinki Declaration. It was approved by the Helsinki University Central Hospital Ethics Committee, and the Regional Committee for Medical and Health Research Ethics South-East Norway. The donors participating in the study have signed written informed consent. Information about patients used in the study can be found in Supplementary Table 1.

**Isolation and culture of human primary T cells, CD34+ HSPCs and fibroblasts**

To isolate human T cells, peripheral blood mononuclear cells (PBMCs) were isolated from human peripheral blood from healthy donors (HD) and patients using Ficoll (StemCell Technologies) gradient centrifugation. Isolated PBMCs were cryopreserved at -150°C. For experiments, PBMCs were thawed and cultured at 1 million cells/mL in ImmunoCult™-XF T Cell Expansion Medium (StemCell Technologies). The basal media was supplemented with 120 U/mL IL-2 (Peprotech), 3 ng/µL IL-7 (Peprotech), 3 ng/µL IL-15 (Peprotech) and 15 µL/mL ImmunoCult™ Human CD3/CD28 T Cell Activator (StemCell Technologies) to make T cell stimulation medium. After incubating cells at 37°C/5% $CO_2$ for three nights, cells were either nucleofected or diluted further with T cell stimulation medium without CD3/CD28 T Cell Activator.

CD34+ hematopoietic stem and progenitor cells (HSPCs) were isolated from cord blood collected during scheduled caesarean sections. Cells were isolated using CD34 MicroBead Kit UltraPure (Miltenyi), and cryopreserved at -150°C. For experiments, CD34+ HSPCs were thawed and cultured at 0.3 million cells/mL in StemSpan™ SFEM II (StemCell Technologies), supplemented with 1X GlutaMax (ThermoScientific), 100 ng/mL human recombinant Flt3-L (Peprotech), 20 ng/mL human recombinant TPO (Peprotech), 100 ng/mL human recombinant SCF (Peprotech), 20 ng/mL human recombinant IL-6 (Peprotech), 1 µM StemRegenin-1 (StemCell Technologies) and 10 µM UM729 (StemCell Technologies) to make HSPC stimulation medium. After incubating cells at 37°C/5% $CO_2$ for three nights, cells were either nucleofected or diluted further with HSPC stimulation medium.

Human fibroblasts were isolated from skin biopsies, expanded in DMEM medium supplemented with low glucose, 1 mM Puryvate and 10% FBS (Gibco) and cryopreserved at -150°C. For experiments, cells were thawed and cultured in same conditions until confluent. Cells were passaged every 3-4 days by dissociating them with TrypLE™ Express Enzyme (Gibco), until cell expansion was complete and cells were nucleofected. All gene editing experiments with fibroblasts were carried out latest at passage 10.

**CRISPR guide RNA (gRNA) design for ADA2, AIRE and RMRP**

We designed 7-18 gRNAs per locus for ADA2, AIRE and RMRP (Supplementary Figure 1), based on available PAM (NGG) sites within the 100bp repair template region centering the mutation site. gRNA sequence information is shown in Table 1, where gRNAs overlapping the mutation sites are marked in red.

**Table 1: gRNA sequences for ADA2, AIRE and RMRP**

| Locus | gRNA | gRNA sequence 5'→3' | gRNA orientation | gRNA length (bp) |
|---|---|---|---|---|
| ADA2 | g#1 | ttccaagtggattctgctgg | FWD | 20 |
| ADA2 | g#2 (WT) | gctggaggattatcggaagc | FWD | 20 |
| ADA2 | g#2 (MUT) | gctggaggattatcagaagc | FWD | 20 |
| ADA2 | g#3 (WT) | tgctggaggattatcggaag | FWD | 20 |
| ADA2 | g#3 (MUT) | tgctggaggattatcagaag | FWD | 20 |
| ADA2 | g#4 | ggattctgctggaggattat | FWD | 20 |
| ADA2 | g#5 | atgttccaagtggattctgc | FWD | 20 |
| ADA2 | g#6 | catcagaaaaatgttccaag | FWD | 20 |
| ADA2 | g#7 | atcctccagcagaatccact | REV | 20 |
| AIRE | g#1 | cagcagtggcccgaagcctc | FWD | 20 |
| AIRE | g#2 (MUT) | gaagcctctggtttgagcca | FWD | 20 |
| AIRE | g#3 (MUT) | aagcctctggtttgagccaa | FWD | 20 |
| AIRE | g#4 (MUT) | gtttgagccaagggagccca | FWD | 20 |
| AIRE | g#5 (MUT) | ggtttgagccaagggagccc | FWD | 20 |
| AIRE | g#6 | aacaaggcccgcagcagcag | FWD | 20 |
| AIRE | g#7 | cttcgggccactgctgctgc | REV | 20 |
| AIRE | g#8 | gcttcgggccactgctgctg | REV | 20 |
| AIRE | g#9 (MUT) | tggctcaaaccagaggcttc | REV | 20 |
| AIRE | g#10 (MUT) | ttggctcaaaccagaggctt | REV | 20 |
| AIRE | g#11 (WT) | gctcccttggctcgaaccag | REV | 20 |

| | | | | |
|---|---|---|---|---|
| AIRE | g#11 (MUT) | gctcccttggctcaaaccag | REV | 20 |
| AIRE | g#12 (MUT) | gctcccttggctcaaaccag | REV | 20 |
| AIRE | g#13 | ggcagcgccctgggctccct | REV | 20 |
| AIRE | g#14 | gcttacgggggcagcgccct | REV | 20 |
| AIRE | g#15 | tgcttacgggggcagcgccc | REV | 20 |
| AIRE | g#16 | ggaaggtcaggtgcttacgg | REV | 20 |
| AIRE | g#17 | gggaaggtcaggtgcttacg | REV | 20 |
| AIRE | g#18 | agggaaggtcaggtgcttac | REV | 20 |
| AIRE | g#19 | cagggaaggtcaggtgctta | REV | 20 |
| RMRP | g#1 | cactctctgcccgaggtccg | FWD | 20 |
| RMRP | g#2 | tgtctacgtgcgtatgcacg | FWD | 20 |
| RMRP | g#3 | cacgtggcactctctgcccg | FWD | 20 |
| RMRP | g#4 | ggcactctctgcccgaggtc | FWD | 20 |
| RMRP | g#5 | gcactctctgcccgaggtcc | FWD | 20 |
| RMRP | g#6 (MUT) | aggtccggggacttcccccct | FWD | 20 |
| RMRP | g#7 (MUT) | tccggggacttccccctagg | FWD | 20 |
| RMRP | g#8 (MUT) | ggacttccccctaggcggaa | FWD | 20 |
| RMRP | g#9 (WT) | gactttcccctaggcggaaa | FWD | 20 |
| RMRP | g#9 (MUT) | gacttccccctaggcggaaa | FWD | 20 |
| RMRP | g#10 (MUT) | acttccccctaggcggaaag | FWD | 20 |
| RMRP | g#11 (MUT) | tccccctaggcggaaagggg | FWD | 20 |
| RMRP | g#12 | gagtcctcagtgtgtagcct | FWD | 20 |
| RMRP | g#13 (MUT) | gggggaagtccccggacctc | REV | 20 |
| RMRP | g#14 (MUT) | agggggaagtccccggacct | REV | 20 |
| RMRP | g#15 (MUT) | tccgcctagggggaagtccc | REV | 20 |
| RMRP | g#16 | ttcctcccctttccgcctag | REV | 20 |

### CRISPR repair template design for ADA2, AIRE and RMRP

Single-stranded DNA repair templates (ssODN) of 100 bp length were designed for ADA2, AIRE and RMRP with +/- 50 bp homology arms surrounding the mutation site. To enable identical editing strategies for ADA2 and AIRE in healthy donors and patients, silent SNPs were added in the designs in addition to mutation correction (WT/MUT→SNP). Four silent SNPs were added for ADA2 and three for AIRE. As RMRP is non-coding, we used non-silent SNPs (WT→SNP, MUT→SNP) for early experiments and mutation correction (MUT→WT) later for functional assessments. ssODN sequence information is presented in Table 2. To further improve homology directed repair in later experiments, we designed asymmetric ssODNs for ADA2, AIRE and RMRP, where we tested 10-40 nt homology arms surrounding the mutation sites. Sequences of asymmetric ssODNs and SNP strategy are presented in Table 2.

#### Table 2: ssODN sequences for ADA2, AIRE and RMRP

| Locus | Repair strategy | ssODN symmetry | ssODN direction (5'→3') | ssODN sequence (100 bp) |
|---|---|---|---|---|
| ADA2 | WT/MUT →SNP | Left 40nt | fwd | cccaagggggatcatgcagttcagatttgctcacccaactccccgtccatcagaaaaatgttccaagtggattctgctggagg actacagaaagcgggtg |
| ADA2 | WT/MUT →SNP | Left 30nt | fwd | atcatgcagttcagatttgctcacccaactccccgtccatcagaaaaatgttccaagtggattctgctggaggactacagaaa gcgggtgcagaacgtca |
| ADA2 | WT/MUT →SNP | Left 20nt | fwd | tcagatttgctcacccaactccccgtccatcagaaaaatgttccaagtggattctgctggaggactacagaaagcgggtgca gaacgtcactgagtttga |
| ADA2 | WT/MUT →SNP | Left 10nt | fwd | tcacccaactccccgtccatcagaaaaatgttccaagtggattctgctggaggactacagaaagcgggtgcagaacgtcact gagtttgatgacaggtga |
| ADA2 | WT/MUT →SNP | Middle | fwd | ccccgtccatcagaaaaatgttccaagtggattctgctggaggactacagaaagcgggtgcagaacgtcactgagtttgatg acaggtgagtagtagttc |
| ADA2 | WT/MUT →SNP | Right 10nt | fwd | cagaaaaatgttccaagtggattctgctggaggactacagaaagcgggtgcagaacgtcactgagtttgatgacaggtgagt agtagttcagaaagcaca |
| ADA2 | WT/MUT →SNP | Right 20nt | fwd | ttccaagtggattctgctggaggactacagaaagcgggtgcagaacgtcactgagtttgatgacaggtgagtagtagttcag aaagcacatgtcccaggc |
| ADA2 | WT/MUT →SNP | Right 30nt | fwd | ttctgctggaggactacagaaagcgggtgcagaacgtcactgagtttgatgacaggtgagtagtagttcagaaagcacatgt cccaggcctgtcatgggg |
| ADA2 | WT/MUT →SNP | Right 40nt | fwd | aggactacagaaagcgggtgcagaacgtcactgagtttgatgacaggtgagtagtagttcagaaagcacatgtcccaggcc tgtcatggggtggcagtgg |
| AIRE | WT/MUT →SNP | Left 40nt | rev | cccttggcacgtaccaaaggcttcgggccactgctgctgcgggccttgttcttcccactgccggagtcttcgaacttgctggga gtgtagaactccccgc |
| AIRE | WT/MUT →SNP | Left 30nt | rev | gccctgggctcccttggcacgtaccaaaggcttcgggccactgctgctgcgggccttgttcttcccactgccggagtcttcgaa cttgctgggagtgtag |
| AIRE | WT/MUT →SNP | Left 20nt | rev | cgggggcagcgccctgggctcccttggcacgtaccaaaggcttcgggccactgctgctgcgggccttgttcttcccactgccg gagtcttcgaacttgct |

| | | | | |
|---|---|---|---|---|
| AIRE | WT/MUT →SNP | Left 10nt | rev | caggtgcttacgggggcagcgccctgggctcccttggcacgtaccaaaggcttcgggccactgctgctgcgggccttgttcttcccactgccggagtctt |
| AIRE | WT/MUT →SNP | Middle | rev | cagggaaggtcaggtgcttacgggggcagcgccctgggctcccttggcacgtaccaaaggcttcgggccactgctgctgcgggccttgttcttcccactg |
| AIRE | WT/MUT →SNP | Right 10nt | rev | ccaggctccccagggaaggtcaggtgcttacgggggcagcgccctgggctcccttggcacgtaccaaaggcttcgggccactgctgctgcgggccttgtt |
| AIRE | WT/MUT →SNP | Right 20nt | rev | gcatcaagagccaggctccccagggaaggtcaggtgcttacgggggcagcgccctgggctcccttggcacgtaccaaaggcttcgggccactgctgctgc |
| AIRE | WT/MUT →SNP | Right 30nt | rev | ggggcgggggcatcaagagccaggctccccagggaaggtcaggtgcttacgggggcagcgccctgggctcccttggcacgtaccaaaggcttcgggcca |
| AIRE | WT/MUT →SNP | Right 40nt | rev | cgctgttcctggggcgggggcatcaagagccaggctccccagggaaggtcaggtgcttacgggggcagcgccctgggctcccttggcacgtaccaaagg |
| RMRP | WT→SNP | Left 40nt | fwd | gatacgcttcttggcggactttggagtgggaagcggggaatgtctacgtgcgtatgcacgtggcactctctgcccgaggtccggggacttccacataggc |
| RMRP | WT→SNP | Left 30nt | fwd | ttggcggactttggagtgggaagcggggaatgtctacgtgcgtatgcacgtggcactctctgcccgaggtccggggacttccacataggcggaaagggga |
| RMRP | WT→SNP | Left 20nt | fwd | ttggagtgggaagcggggaatgtctacgtgcgtatgcacgtggcactctctgcccgaggtccggggacttccacataggcggaaaggggaggaacagagt |
| RMRP | WT→SNP | Left 10nt | fwd | aagcggggaatgtctacgtgcgtatgcacgtggcactctctgcccgaggtccggggacttccacataggcggaaaggggaggaacagagtcctcagtgtg |
| RMRP | WT→SNP | Middle | fwd | tgtctacgtgcgtatgcacgtggcactctctgcccgaggtccggggacttccacataggcggaaaggggaggaacagagtcctcagtgtgtagcctagga |
| RMRP | WT→SNP | Right 10nt | fwd | cgtatgcacgtggcactctctgcccgaggtccggggacttccacataggcggaaaggggaggaacagagtcctcagtgtgtagcctaggatacaggcctt |
| RMRP | WT→SNP | Right 20nt | fwd | tggcactctctgcccgaggtccggggacttccacataggcggaaaggggaggaacagagtcctcagtgtgtagcctaggatacaggccttcagcacgaac |
| RMRP | WT→SNP | Right 30nt | fwd | tgcccgaggtccggggacttccacataggcggaaaggggaggaacagagtcctcagtgtgtagcctaggatacaggccttcagcacgaaccacgtcctca |
| RMRP | WT→SNP | Right 40nt | fwd | ccggggacttccacataggcggaaaggggaggaacagagtcctcagtgtgtagcctaggatacaggccttcagcacgaaccacgtcctcagcttcacaga |
| RMRP | MUT→SNP | Middle | fwd | tgtctacgtgcgtatgcacgtggcactctctgcccgaggtccggggactttcacataggcggaaaggggaggaacagagtcctcagtgtgtagcctagga |
| RMRP | MUT→WT | Middle | fwd | tgtctacgtgcgtatgcacgtggcactctctgcccgaggtccggggactttcccctaggcggaaaggggaggaacagagtcctcagtgtgtagcctagga |

### Nucleofection of human primary T cells, CD34+ HSPCs and fibroblasts

Human T cells, CD34+ HSPCs and fibroblasts were nucleofected using 4-D Nucleofector system and 96-well unit (Lonza). gRNAs were made by annealing crRNA (IDT) and trcrRNA (IDT) as according to the manufacturer's instructions. RNPs were prepared by mixing 61pmol Alt-R™ S.p. Cas9 Nuclease V3 (IDT) with 100 pmol annealed gRNA per sample and incubating at 37°C for 15 min, after which 100 pmol ssODN (IDT) was added.

For nucleofection, 0.5 or 1 million T cells, 0.3 million HSPCs and 1 million fibroblasts per sample were resuspended in 20 uL Primary P3 electroporation buffer (Lonza) and mixed carefully with the RNPs. Cells were nucleofected with the following programs: EO-115 (T cells), DZ-100 (HSPCs), CA-137 (fibroblasts).

After nucleofection of T cells, 85 µL T cell recovery medium (basal medium supplemented with 250U/mL IL-2) was added into the electroporation plate, followed by 15 min incubation at 37°C/5% $CO_2$. Afterwards, cells were transferred into 24-well (1 million cells/sample) or 48-well (0.5 million cells/sample) plates to grow and incubated at 37°C/5% $CO_2$. Cells were split 1:1 or as necessary with T cell recovery medium 24h and 72h after nucleofection. Cells were collected for downstream analyses 4 days, or alternatively 6-8 days, after nucleofection depending on the experiment.

After nucleofection of HSPCs, 85 µL HSPC stimulation medium (basal medium supplemented with aforementioned cytokines) was added into the electroporation plate, followed by 15 min incubation at 37°C/5% $CO_2$. Afterwards, cells were transferred into 48-well culture plates to grow and incubated at 37°C/5% $CO_2$. HSPC stimulation medium was added 24 and 72h after nucleofection if necessary. Cells were collected for downstream analyses 4 days after nucleofection.

After nucleofection of fibroblasts, 85 µL fibroblast culture medium (basal medium supplemented with aforementioned reagents) was added into the electroporation plate, followed by 15 min incubation at 37°C/5% $CO_2$. Afterwards, cells were transferred into 6-well culture plates to grow and incubated at 37°C/5% $CO_2$. Medium was changed 24h after nucleofection, and samples were trypsinized and collected for downstream analyses 4 days after nucleofection.

**Flow cytometry**

**1. Characterization of immune cells in HD PBMCs**

PBMC samples from day 1, 4 and 8 of the pipeline were prepared for flow cytometry analysis by washing 0.5 million cells per sample once with RT PBS, followed by blocking for 10 min at RT with 10% human serum in PBS. Cells were then stained in the dark for 30 min at 4°C by adding 50 µL antibody cocktail per sample, as described in Table 3. After staining, cells were washed twice with 200 µL flow buffer (eBioscience) and resuspended in 250 µL flow buffer. Samples were stored in the dark at 4°C until flow cytometry. The flow analyses were performed on LSRII (BD Bioscience) at the Flow Cytometry Core Facility at Oslo University Hospital (Oslo, Norway). Data analysis was done with FlowJo software (FlowJo LLC, Ashland, OR).

**Table 3: Markers used for immune cell characterization**

| Marker | Color | Clone | Vendor | Catalog | Dilution |
|---|---|---|---|---|---|
| CD14 | PerCP-Cy5.5 | 61D3 | eBioscoence | 45-0149-42 | 1:200 |
| CD15 | FITC | 3G8 | Biolegend | 302001 | 1:200 |
| CD56 | FITC | NCAM16.2 | BD | 664524 | 1:200 |
| CD4 | AlexaFluor700 | RPA-T4 | BD Pharmingen | 557922 | 1:200 |
| CD3 | BV421 | UCHT1 | BD Horizon | 562426 | 1:200 |
| CD20 | BV786 | 2H7 | Biolegend | 302356 | 1:100 |
| CD8 | PE | 4B9 | eBioscience | 12-0087-42 | 1:100 |
| LiveDead near IR | | | Thermo Fisher | L34992 | 1:1000 |

### 2. CD4+ CD8+ T cell sorting panel for fluorescence-activated cell sorting

T cells from DADA2 patient and HD were collected on day 8 of the pipeline and prepared for flow cytometry analysis by collecting 2 million cells per sample and washing them once with ice-cold PBS. Cells were resuspended with 200 µL Live/Dead dye combined FcR Blocking Reagent (Miltenyi) and samples were stained in the dark at 4°C for 30 min. Cells were then stained in the dark for 30 min at 4°C by adding 50 µL antibody cocktail per sample, as described in Table 4. After staining, cells were washed once in cold flow buffer (eBioscience) and resuspended in cold flow buffer, followed by FACS (SONY SH800S) at Centre for Molecular Medicine Norway at University of Oslo, Norway. Data analysis was done with FlowJo software (FlowJo LLC, Ashland, OR).

**Table 4: Markers used for CD4+ and CD8+ T cell sorting panel**

| Marker | Color | Clone | Vendor | Catalog | Dilution |
|---|---|---|---|---|---|
| CD19 | PE | HIB19 | BioLegend | 302207 | 1:100 |
| CD14 | PE | HCD14 | BioLegend | 325605 | 1:100 |
| CD11c | PE | 3.9 | BioLegend | 301605 | 1:100 |
| CD56 | PE | HCD56 | BioLegend | 318305 | 1:100 |
| CD3 | Pacific Blue | SK7 | BioLegend | 344823 | 1:50 |
| CD4 | APC-Cy7 | OKT4 | BioLegend | 317417 | 1:50 |
| CD8a | PerCP-Cy5.5 | RPA-T8 | BioLegend | 301031 | 1:100 |
| LIVE/DEAD Fixable Red | | | Invitrogen | L34971 | 1:500 |

### 3. T cell proliferation assay in CHH patients

CHH patient T cells from day 20 of the pipeline were collected and washed once with PBS. Cells were resuspended in PBS at 2 million cells/mL. CFSE working solution (2 µM) was prepared right before staining from CellTrace™ CFSE Cell Proliferation Kit (Invitrogen), where stock (5mM) was first diluted with PBS. To stain cells, equal volume of CFSE working solution and PBS were added to get a final concentration of 1 uM CFSE. Cells were immediately vortexed for 10 s, followed by incubation in the dark at 37°C, 5% CO2 for 5 min, including a brief vortexing step at 2.5 min of incubation. Immediately after incubation, equal volume of cold human serum (Sigma) was added on cells. Cells were centrifuged, washed twice with PBS and resuspended in Immunocult medium at 4 million cells/mL. Cell suspension (50 uL per sample) was added per well on a 96-well U bottom plate (Thermo Fisher) to get a final concentration of 0.2 million cells/mL per well with a final concentration of 250 U/mL IL-2 (Peprotech). After staining, cells were incubated for four days at 37°C, 5% $CO_2$. On day 24 of the pipeline, cells were stained for flow cytometry. Samples were washed once with PBS and resuspended in 50 uL Live/Dead staining with Fc blocking reagent (Miltenyi) per sample and incubated in the dark for 30 min at 4°C. Samples were centrifuged and resuspended in 50 uL antibody cocktail as described in Table 5 and stained in the dark for 30 min at 4°C. Afterwards, cells were washed two times with flow buffer (eBioscience) and resuspended in flow buffer for flow cytometry analysis. The flow analyses were performed on LSRII (BD Bioscience) at the Flow Cytometry Core Facility at Oslo University Hospital (Oslo, Norway). Data analysis was done with FlowJo software (FlowJo LLC, Ashland, OR).

**Table 5: Antibodies and reagents used for CFSE T cell proliferation assay**

| Marker | Color | Clone | Vendor | Catalog | Dilution |
|---|---|---|---|---|---|
| CFSE | CFSE | | | C34554A | 1µM |
| CD19 | Pacific Blue | HIB19 | Biolegend | 302232 | 1:200 |
| CD14 | Pacific Blue | M5E2 | Biolegend | 301828 | 1:200 |
| CD56 | Pacific Blue | MEM-188 | Biolegend | 304629 | 1:200 |
| CD11c | V450 | B-ly6 | BD Biosciences | 560369 | 1:200 |
| Live/Dead Fixable Violet | | | Invitrogen | L34963 | 1:500 |
| CD8a | Alexa Fluor 594 | RPA-T8 | Biolegend | 301056 | 1:100 |
| CD4 | APC-Cy7 | OKT4 | BioLegend | 317417 | 1:100 |

### Assessment of *in silico* gRNA design tools

To evaluate predictive power of available *in silico* gRNA design tools against *in vitro* gRNA screening data, we selected the following tools: Atum (https://www.atum.bio/eCommerce/cas9/input), Benchling (benchling.com), CHOPCHOP (chopchop.cbu.uib.no), CRISPOR (crispor.tefor.net/), DeepSpCas9 (deepcrispr.info/DeepSpCas9/), EuPaGDT (grna.ctegd.uga.edu/) and IDT gRNA design tool (eu.idtdna.com/site/order/designtool/index/CRISPR_SEQUENCE). We used 100 bp mutant-specific sequences with 50 bp homology arms from the mutation site as input (target sequence) for the tools. Three gRNAs with highest predicted efficiency were chosen from the tool output and compared against three best *in vitro* validated gRNAs from patient T cells.

**On-target editing assessment by Droplet Digital PCR (ddPCR)**

ddPCR assays were performed to assess HDR and NHEJ editing for ADA2, AIRE, Enh4-1, CTCF1, RNF2 and RMRP. We used previously published ddPCR oligos for Enh4-1, CTCF1, RNF2[1] and designed new ddPCR oligos for ADA2, AIRE and RMRP, described in Table 6. ddPCR was performed using the QX200 system (Bio-Rad) as previously described[1]. In short, 8 µl of DNA (concentration normalized to 8 ng/µl), primers (900 nM), reference probe (250 nM), and HDR or NHEJ probe (250 nM). The HDR and NHEJ detection occurred in two separate ddPCR reactions. Each reaction was then loaded into a sample well of an eight-well disposable cartridge (DG8; Bio-Rad Laboratories) along with 70 µl of droplet generation oil (Bio-Rad Laboratories). Droplets were formed using a QX200 Droplet Generator (Bio-Rad Laboratories). Droplets were transferred to a 96-well PCR plate, heat-sealed with foil, and amplified using a conventional thermal cycler. The thermocycling protocol was the following: (1) 95°C - 10 min, (2) 94°C – 30 s, 56°C – 3 min, step repeated 42 times (3) 98°C – 10 min, (4) 4°C – hold. The resulting PCR products were loaded on a QX200 Droplet Reader (Bio-Rad Laboratories), and the data was analyzed using QuantaSoft software (Bio-Rad Laboratories).

**Table 6: ddPCR oligos for ADA2, AIRE and RMRP**

| Target gene | ddPCR primer fwd | ddPCR primer rev | ddPCR probe reference | ddPCR probe HDR | ddPCR probe NHEJ |
|---|---|---|---|---|---|
| ADA2 | GGTGAGG AATGTCA CCTACA | GTACCA AGGGA GACACC TAC | **WT&MUT:**<br><br>/5'FAM/GCCACATCTGT TTCACCCCA/3'BHQ_1/ | **WT/MUT→SNP:**<br>/5'HEX/CTGGAGGACTAC AGAAAGCGG/3'BHQ_1/ | **WT→SNP:**<br>/5'HEX/ATTATCGGAAGC GGGTGCAGA/3'BHQ_1/<br><br>**MUT→SNP:**<br>/5'HEX/ATTATCAGAAGC GGGTGCAGA/3'BHQ_1/ |

| | | | | | |
|---|---|---|---|---|---|
| AIRE | TCTACACTCCCAGCAAGTTC | GGAAGGTCAGGTGCTTACG | **WT&MUT:**<br><br>/5'FAM/TCCGGCAGTGGGAAGAACAA/3'BHQ_1/ | **WT/MUT→SNP:**<br>/5'HEX/AAGCCTTTGGTACGTGCCAAG/3'BHQ_1/ | **WT→SNP:**<br>/5'HEX/CGAAGCCTCTGGTTCGAGC/3'BHQ_1/<br><br>**MUT→SNP:**<br>/5'HEX/CGAAGCCTCTGGTTTGAGC/3'BHQ_1/ |
| RMRP | GCTTCTTGGCGGACTTTG | ATACTACTCTGTGAAGCTGAGG | **WT&MUT:**<br><br>/5'FAM/TGGGAAGCGGGGAATGTCTA/3'BHQ_1/ | **WT→SNP:**<br>/5'HEX/GGACTTCCACATAGGCGGAA/3'BHQ_1/<br><br>**MUT→SNP:**<br>/5'HEX/GGACTTTCACATAGGCGGAA/3'BHQ_1/ | **WT→SNP:**<br>/5'HEX/ACTTTCCCCTAGGCGGAAAG/3'BHQ_1/<br><br>**MUT→SNP:**<br>/5'HEX/ACTTCCCCCTAGGCGGAAAG/3'BHQ_1/ |

**On-target editing assessment by amplicon sequencing**

Amplicon sequencing libraries for assessing on-target editing for *ADA2*, *AIRE* and *RMRP* were prepared from gDNA samples as previously described[1]. In short, library preparation was performed using a two-step PCR method. For the first PCR, a pair of target-specific primers were designed to amplify a 150 bp area surrounding the cutting site. Each target primer additionally includes an extension at the 5' end: for forward primers, this contains the Illumina Read1 primer sequence (see below, nucleotides in bold) and an 8 bp UMI (nucleotides underlined), and for the reverse primers, this contains the Illumina Read2 primer sequence only (see below, nucleotides in bold):

ADA2 fwd 5'→3': **ACACTCTTTCCCTACACGACGCTCTTCCGATCT**NNNNNNNNTCATGCAGTTCAGATTTGCTCAC
ADA2 rev 5'→3': **GTGACTGGAGTTCAGACGTGTGCTCTTCCGATCT**GGCCTGGGACATGTGCTTTC

AIRE fwd 5'→3': **ACACTCTTTCCCTACACGACGCTCTTCCGATCT**NNNNNNNNactcccagcaagttcgaaga
AIRE rev 5'→3': **GTGACTGGAGTTCAGACGTGTGCTCTTCCGATCT**GGGGGCATCAAGAGCCAG

RMRP fwd 5'→3': **ACACTCTTTCCCTACACGACGCTCTTCCGATCT**NNNNNNNNgagtgggaagcggggaatg
RMRP rev 5'→3': **GTGACTGGAGTTCAGACGTGTGCTCTTCCGATCT**AGCTGAGGACGTGGTTCGT

First PCR was performed with reagents listed in Table 7. The thermocycling protocol was the following: (1) 98°C - 30 s, (2) 98°C – 10 s, 57°C (ADA2, RMRP)/58°C (AIRE) – 10 s, 72°C – 20 s, step repeated 30 times (3) 72°C – 5 min, (4) 4°C – hold.

**Table 7: Amplicon seq first PCR components**

| Reagent | Vendor | Final concentration |
|---|---|---|
| Nuclease-free water | Ambion | Add up to 20 |
| 5× Phusion GC Buffer | Thermo Fisher Scientific | 1X |
| 10 mM dNTPs | Thermo Fisher Scientific | 200 µM |

| | | |
|---|---|---|
| 10 µM Primer fwd | IDT | 0.5 µM |
| 10 µM Primer rev | IDT | 0.5 µM |
| Betaine | Sigma Aldrich | 1 M |
| Phusion Hot Start II DNA Polymerase | Thermo Fisher Scientific | 0.02 U/µl |
| Template DNA | N/A | 100 ng |

For the second PCR, the amplified products were purified using AMPure XP magnetic (Beckman Coulter, #A63882) according to the manufacturer’s instructions, pooled and annealed with i5 and i7 Illumina Index primers (Table 8). Both primers contain flow-cell-binding region (highlighted in bold), index region (underlined) and Illumina Read1 or Read2 primer binding regions, correspondingly (italics).

**Table 8: Amplicon sequencing second PCR primers**

| | |
|---|---|
| i5-PCR Index 9 | **AAT GAT ACG GCG ACC ACC GAG ATC TA**TTGCTTGC_AC ACT CTT TCC CTA CAC GAC GCT CTT CCG ATC* T_ |
| i5-PCR Index 10 | **AAT GAT ACG GCG ACC ACC GAG ATC TA**GAGAGGTT_AC ACT CTT TCC CTA CAC GAC GCT CTT CCG ATC* T_ |
| i5-PCR Index 11 | **AAT GAT ACG GCG ACC ACC GAG ATC TA**ACCTGGTT_AC ACT CTT TCC CTA CAC GAC GCT CTT CCG ATC* T_ |
| i5-PCR Index 13 | **AAT GAT ACG GCG ACC ACC GAG ATC TA**CGGAACAA_AC ACT CTT TCC CTA CAC GAC GCT CTT CCG ATC* T_ |
| i5-PCR Index A505 | **AAT GAT ACG GCG ACC ACC GAG ATC TA**CTAATCGA_AC ACT CTT TCC CTA CAC GAC GCT CTT CCG ATC* T_ |
| i5-PCR Index A506 | **AAT GAT ACG GCG ACC ACC GAG ATC TA**CTAGAACA_AC ACT CTT TCC CTA CAC GAC GCT CTT CCG ATC* T_ |
| i5-PCR Index A507 | **AAT GAT ACG GCG ACC ACC GAG ATC TA**TAAGTTCC_AC ACT CTT TCC CTA CAC GAC GCT CTT CCG ATC* T_ |
| i5-PCR Index A508 | **AAT GAT ACG GCG ACC ACC GAG ATC TA**TAGACCTA_AC ACT CTT TCC CTA CAC GAC GCT CTT CCG ATC* T_ |
| i7-PCR Index 13 | **CAA GCA GAA GAC GGC ATA CGA GAT**TTCCTCCT_G TGA CTG GAG TTC AGA CGT GTG CTC TTC CGA TC*T_ |
| i7-PCR Index 14 | **CAA GCA GAA GAC GGC ATA CGA GAT**TGCTTGCT_G TGA CTG GAG TTC AGA CGT GTG CTC TTC CGA TC*T_ |
| i7-PCR Index 15 | **CAA GCA GAA GAC GGC ATA CGA GAT**GGTGATGA_G TGA CTG GAG TTC AGA CGT GTG CTC TTC CGA TC*T_ |
| i7-PCR Index 16 | **CAA GCA GAA GAC GGC ATA CGA GAT**AACCTACG_G TGA CTG GAG TTC AGA CGT GTG CTC TTC CGA TC*T_ |
| i7-PCR Index A705 | **CAA GCA GAA GAC GGC ATA CGA GAT**ACCCAGCA_G TGA CTG GAG TTC AGA CGT GTG CTC TTC CGA TC*T_ |
| i7-PCR Index A706 | **CAA GCA GAA GAC GGC ATA CGA GAT**AACCCCTC_G TGA CTG GAG TTC AGA CGT GTG CTC TTC CGA TC*T_ |
| i7-PCR Index A707 | **CAA GCA GAA GAC GGC ATA CGA GAT**CCCAACCT_G TGA CTG GAG TTC AGA CGT GTG CTC TTC CGA TC*T_ |

| | |
|---|---|
| i7-PCR Index A708 | **CAA GCA GAA GAC GGC ATA CGA GAT**CACCACAC_G TGA CTG GAG TTC AGA CGT GTG CTC TTC CGA TC*T_ |
| i7-PCR Index A709 | **CAA GCA GAA GAC GGC ATA CGA GAT**GAAACCCA_G TGA CTG GAG TTC AGA CGT GTG CTC TTC CGA TC*T_ |
| i7-PCR Index A710 | **CAA GCA GAA GAC GGC ATA CGA GAT**TGTGACCA_G TGA CTG GAG TTC AGA CGT GTG CTC TTC CGA TC*T_ |
| i7-PCR Index A711 | **CAA GCA GAA GAC GGC ATA CGA GAT**AGGGTCAA_G TGA CTG GAG TTC AGA CGT GTG CTC TTC CGA TC*T_ |
| i7-PCR Index A712 | **CAA GCA GAA GAC GGC ATA CGA GAT**AGGAGTGG_G TGA CTG GAG TTC AGA CGT GTG CTC TTC CGA TC*T_ |

Second PCR was performed with reagents listed in Table 9. The thermocycling protocol was the following: (1) 98°C - 30 s, (2) 98°C – 10 s, 58°C – 10 s, 72°C – 20 s, step repeated 10 times (3) 72°C – 5 min, (4) 4°C – hold.

**Table 9: Amplicon seq second PCR components**

| Reagent | Vendor | Final concentration |
|---|---|---|
| Nuclease-free water | Ambion | Add up to 20 |
| 5× Phusion GC Buffer | Thermo Fisher Scientific | 1X |
| 10 mM dNTPs | Thermo Fisher Scientific | 200 µM |
| 10 µM Primer fwd | IDT | 0.25 µM |
| 10 µM Primer rev | IDT | 0.25 µM |
| Betaine | Sigma Aldrich | 1 M |
| Phusion Hot Start II DNA Polymerase | Thermo Fisher Scientific | 0.02 U/µl |
| Template DNA | N/A | 5 ng |

PCR products were purified using AMPure XP magnetic beads (Beckman Coulter) according to the manufacturer's instructions and DNA concentrations were measured with Qubit HS kit (Thermo Fisher Scientific). Final sample libraries were sequenced using Illumine MiSeq v2 Micro flow cell, including 10% PhiX. Data analysis was performed using the ampliCan software package[2].

**Off-target assessment by Genome-wide, unbiased identification of DSBs enabled by sequencing (GUIDE-seq)**

One million patient and healthy donor T cells/sample were nucleofected on day 5 of the pipeline, as previously described, with RNPs containing selected gRNAs 100 pmol, Cas9 nuclease at 61 pmol and dsODN at 30 pmol/sample. Cells were transferred into 24w plates after nucleofection with 500 uL T cell recovery medium and split 1:1 with T cell recovery medium 24h and 72h after nucleofection. Samples were collected for GUIDE-seq sample processing and ddPCR 4 days after nucleofection.

The blunt-ended dsODN used in our GUIDE-Seq experiments was the same as which was used in the original publication [3] and then was prepared by annealing the two modified oligonucleotides of the following compositions:

5'- P-G*T**TTAATTGAGTTGTCATATGTTAATAACGGT**A*T -3' and
5'- P-A*T**ACCGTTATTAACATATGACAACTCAATTAA**A*C -3'
P represents a 5' phosphorylation and * indicates a phosphorothioate linkage.

The GUIDEseq protocol [3] was adapted from certain modifications described below. Briefly, gDNA was sheared with a Bioruptor® Pico Sonication System (Diagenode) to an average length of 500 bp. End-repair was done with Fast DNA End Repair Kit (Thermo Fisher Scientific), A-tailing with Taq DNA Polymerase, native (Thermo Fisher Scientific) and ligation of half-functional adapters, incorporating 8-nt random molecular index was done using T4 DNA Ligase (Thermo Fisher Scientific), all according to the manufacturer's instructions. Between each step, DNA was purified using AMPure XP SPRI beads (Backman Coulter) and eluted with TE buffer, pH 8.0 (Invitrogen). Two rounds of nested anchored PCR, with primers complementary to the oligo tag, were used for target enrichment. Before library pooling, the quality of the final products was tested using High Sensitivity DNA Kit (Agilent) on Bioanalyzer 2100 (Agilent) according to the manufacturer instructions and NanoDrop (Thermo Fisher Scientific). Based on the average size estimated from Bioanalyzer, equal number of particles from each sample were pulled to achieve the final volume 20µl containing $1{,}2 \times 10^{10}$.

Sample was delivered together with custom sequencing primer Index-1 and Read-2 and sequenced at The Department of Core Facilities, Oslo University Hospital, Norway. Denaturated library was loaded onto the Miseq according to Illumina's standard protocol for sequencing with an Illumina Miseq Reagent Kit V2 - 300 cycle (2 x 150 bp paired end).

Data analysis was performed following the GUIDE-Seq analysis pipeline from Zhu et al. 2017, but adjusted for allowing bulges between sgRNA and off-target sites with editing distance of 4. We used custom scripts (https://git.app.uib.no/valenlab/t_cell_editing_pipeline/) with cutadapt v2.8 (TTGAGTTGTCATATGTTAATAACGGTAT and ACATATGACAACTCAATTAAAC). Afterward, the data was aligned to the human genome (hg38v34) using bwa v0.7.17-r1188. CHOPOFF (https://github.com/JokingHero/CHOPOFF.jl) was used to find all off-target sites with edit distance up to 4, allowing for mismatches, deletions and insertions. Final off-targets were normalized against control data (transfected with dsODN only). Control data was processed in the same pipeline as the modified Cas9 samples.

**Cas9WT and Cas9-SNAP *in vitro* mRNA transcription (IVT)**

Cas9WT and Cas9-SNAP mRNA were prepared with HiScribe T7 ARCA mRNA Kit with tailing (NEB-Bionordika), according to the manufacturer's instructions. Total of 8000 ng stock plasmid was digested with 2 µl FastDigest MssI enzyme (Thermo Fisher Scientific) in the supplemented restriction-digestion buffer with a total reaction volume of 20 µl. Incubation was carried out at 37°C overnight. Length of the digested product was confirmed by gel electrophoresis. For the IVT reaction, 1000 ng of the linearized plasmid was mixed with 10 µl of 2xARCA/NTP mix and 2 µl of T7 RNA Polymerase mix, and the reaction was incubated for 30 min at 37°C. Sequentially, 2 µl of DNAse enzyme was added and the mixture was incubated at 37°C for 15 min. Poly(A) tailing step was performed by adding 20 µl of milliQ (RNAse free), 5 µl of 10× PolyA polymerase reaction buffer, and 5 µl of 10× PolyA polymerase directly to the IVT reaction, which was incubation at 37°C for 30 min. mRNA was purified using LiCl solution, as described in the manufacturer's protocol. Aliquots were frozen in –80°C for later use.

**Synthesis of $O^6$-Benzylguanine coupled repair templates**

BG-coupled repair template oligos for ADA2 and AIRE were prepared as previously described [4, 5]. In short, a coupling reaction of BG-GLA-NHS (New England BioLabs) and NH2-oligo (IDT) in HEPES buffer pH 8.5 (Invitrogen) was performed. Following coupling reactions, the BG-oligos were purified with ethanol precipitation as and stored at -20°C until later use.

**Cas9-SNAP nuclease production**

To test the editing performance of BG-coupled repair oligo in combination with the Cas9-SNAP fusion protein, the Cas9-SNAP protein was produced as protein. The pTH24-Cas9-SNAP construct was transformed into *E. coli* BL21(DE3) T1R cells and cultivated in Terrific Broth (TB) medium. Protein expression was induced with isopropyl-D-1-thiogalactopyranoside, and protein purified by immobilized metal-ion chromatography, followed by size exclusion chromatography (SEC). The purified protein was stored in 20 mM HEPES supplemented with 300 mM NaCl, 10% glycerol and 2 mM TCEP to pH 7.5. Aliquots were flash-frozen in liquid nitrogen and stored at -80℃ until experiments.

**HDR enhancing compound screen in healthy donor T cells**

We selected 33 previously published HDR enhancing compounds for the screen, described in Table 10. Compounds were dissolved in DMSO and each of them were assessed at three concentrations in HD T cells against DMSO vehicle control. As described previously, 0.5 million T cells/sample were nucleofected, transferred to 48-w cell culture plates and incubated in T cell recovery medium containing the compounds for 24h. ells were split 1:1 in T cell recovery medium without the compound, 24h and 72h after nucleofection. Samples were collected for gDNA extraction and ddPCR 96h after nucleofection.

**Table 10: List of HDR enhancing compounds and tested concentrations**

| Compound | Conc 1 (µM) | Conc 2 (µM) | Conc 3 (µM) |
|---|---|---|---|
| ABT263 [6] | 0,25 | 0,5 | 1 |
| AICAR[7] | 10 | 20 | 40 |
| B02 [7] | 10 | 20 | 40 |
| Brefeldin A[8] | 0,05 | 0,1 | 0,2 |
| Entinostat[9] | 2,5 | 5 | 10 |
| EPZ5676[10] | 0,05 | 0,1 | 0,2 |
| IC86621[11] | 100 | 200 | 400 |
| KU0060648[12, 13] | 0,125 | 0,25 | 0,5 |
| KU55933[14, 15] | 1,5 | 3 | 6 |
| L755507[8] | 2,5 | 5 | 10 |
| Mirin[14] | 1,5 | 3 | 6 |
| MLN4924[7] | 0,25 | 0,5 | 1 |
| M3814[16] | 1 | 2 | 4 |
| Nexturastat A[17] | 1,25 | 2,5 | 5 |
| NSC 15520[7] | 2,5 | 5 | 10 |
| NSC 19630[7] | 0,5 | 1 | 2 |
| NU7026[7] | 10 | 20 | 40 |
| NU7441[13] | 1 | 2 | 4 |
| Panobinostat[9] | 0,05 | 0,1 | 0,2 |
| PFM01[14] | 5 | 10 | 20 |
| Resveratrol[18] | 0,5 | 1 | 25 |
| Ricolinostat[17] | 1,25 | 2,5 | 5 |

| | | | |
|---|---|---|---|
| Romidepsin[19, 20] | 0,01 | 0,025 | 0,1 |
| RS-1[21] | 5 | 10 | 20 |
| Rucaparib[22, 23] | 2,5 | 5 | 10 |
| SCR7 pyrazine | 2,5 | 1 | 5 |
| STL127705[7, 24] | 2,5 | 5 | 10 |
| TDRL-505[14] | 10 | 20 | 40 |
| Trichostatin A[7, 25] | 0,005 | 0,01 | 0.1 |
| Valproic acid[20, 26] | 5 | 10 | 20 |
| Wortmannin[27] | 0,01 | 0,02 | 0,04 |
| Crispy mix[7]* | * | Not tested | Not tested |
| IDT ALT-R enhancer V2 | 1 | Not tested | Not tested |

*20 µM NU7026, 0.01 µM Trichostatin A, 0.5 µM MLN4924, 5uM NCS15520

**Validating cell cycle inhibitors in healthy donor T cells**

We selected 10 previously published HDR enhancing cell cycle inhibitors for validating in HD T cells and assessed at three concentrations against DMSO vehicle control, as described in Table 11. Cells were either pre-treated with the compounds or vehicle for 24h before nucleofection, followed by nucleofection and incubation without compounds, or treated for 24h after nucleofection. For both conditions, 0.5 million cells per sample were nucleofected. For both groups, cells were split 1:1 in T cell recovery medium without compounds 24h and 72h after nucleofection. Samples were collected for gDNA extraction and ddPCR 96h after nucleofection.

**Table 11: List of cell cycle inhibitors and tested concentrations**

| Compound | Conc 1 (µM) | Conc 2 (µM) | Conc 3 (µM) |
|---|---:|---:|---:|
| ABT-751[28] | 0,175* | 0,35* | 0,7* |
| Aphidicolin[29] | 1* | 2* | 4* |
| AZD7762[30] | 0,5 | 1 | 2 |
| Hydroxy urea[29] | 62,5 | 125 | 250 |
| Lovastatin[29] | 20 | 40 | 80 |

| | | | |
|---|---|---|---|
| Mimosine[29] | 100 | 200 | 400 |
| Nocodazole[29] | 0,1* | 0,2* | 0,4* |
| PHA-767491[31] | 5 | 10 | 20 |
| Thymidine[29] | 1250 | 2500 | 5000 |
| XL413[31] | 5 | 10 | 20 |

* Concentration reported as µg/mL instead of µM

**PacBio sequencing and variant calling of CRISPR edited healthy donor T cells**

T cells from a healthy donor with written consent for sequencing were cultured and edited as previously described. The cells were either unedited and treated with 0.5 µM KU0060648 or DMSO or ADA2-edited and treated with 0.5 µM KU0060648 or DMSO. Cells were collected six days after editing on day 10 of the pipeline and DNA was extracted from 5 million cells per sample using Blood & Cell Culture DNA Kits (Qiagen). All samples were extracted according to the manufacturer's instructions for Cell cultures described in "QIAGEN® Genomic DNA Handbook, June 2015". The concentration, purity and size of the DNA was estimated using both NanoDrop (Thermo Fisher Scientific) and a Qubit fluorometer (Invitrogen) and checked on the agarose gel (0.5%, 35V, 16-18 hours runtime) containing 500 ng of each sample, with an appropriate ladder as a reference standard: Quick-Load 1 kb Extend DNA Ladder (New England Biolabs).

Library preparations for PacBio HiFi sequencing were done by the Norwegian sequencing Centre on 8M SMRT cells using Revio HiFi prep kit and Sequencing chemistry v2.0. The sequencing data was demultiplexed with the Demultiplexing pipeline on SMRT Link v10.2.0.1333434. Circular consensus sequencing (CCS) reads were then generated for demultiplexed polymerase reads and further demultiplexed using the barcoded primer sequences. The HiFi sequencing reads were separated and indexed with the provided barcode ID.

The HiFi sequencing reads were aligned with pbmm2 v1.13.0 with options "--preset HIFI --bam-index BAI --sort". Structural variants were called with pbsv v2.9.0, small variants with deepVariant v1.6.0. All possible mismatches, deletions and insertions were extracted from aligned reads using custom scripts (https://git.app.uib.no/valenlab/t_cell_editing_pipeline/-/tree/main/katariina_pacbio). We normalized data using two control samples, and focused on sites that were potential sgRNA off-target within distance of 4, allowing for bulges. Additionally, transversion ratio plot and codon signature analysis showed no global effects of CRISPR activity.

**CellTiter-Glo cell viability assay for HDR enhancing compound toxicity assessment**

HDR enhancing compound toxicity was assessed by CellTiter-Glo viability assay (Promega) according to manufacturer's instructions. In short, 50 µL of T cell suspension per sample was transferred into while opaque 96-w plates (Thermo Fisher), followed by adding 100 µL RT CellTiter-Glo assay buffer per sample. Plate was covered with aluminum foil and placed on a plate shaker at 500 rpm for 5 min. Afterwards, plate was incubated for 10 min at RT while still covered. After incubation, foil was removed and luminescence values from the plate were assessed by BioTek Synergy Neo2 Instrument (Agilent). To analyze results, background values from medium alone were subtracted from sample values and data was analyzed according to manufacturer's instructions.

**scRNAseq in HD and DADA2 patient T cells**

**1. Cell culture and processing**

Cells were cultured and nucleofected as previously described and sorted with FACS on day 8 of the pipeline into 384 well plates containing 2 uL of lysis buffer[$H_2O$: 1.31 uL, RNase Inhibitor 0.05 uL, ERCC (1:30000) 0.05 uL, 10% Triton (0.04 uL), 10 mM dNTP (0.5 uL) and 100 uM oligo dT (0.05 uL)]. After sorting, the plates were spun down at 2000g, 4°C for 5 min and then the plate was snap frozen on dry ice and kept at -80 until further processing. Reagents for sample processing are described in Table 12.

**Table 12: Reagents for scRNAseq**

| Reagent | Vendor |
|---|---|
| Maxima H Minus Reverse Transcriptase | Thermo Fisher |
| psfTn5 | Addgene |
| KAPA HiFi HotStart ReadyMix | Roche |
| Lambda Exonuclease | BioNordika |
| Tween-20 | Sigma Aldrich |
| 10% SDS solution | Teknova |
| Magnesium Chloride (1 M) | Sigma Aldrich |
| Triton X-100 | Sigma Aldrich |
| KAPA HiFi PCR kit with dNTPs | Roche |
| Betaine (5 M) | Sigma Aldrich |
| UltraPure DNase/RNase Free Distilled Water | Thermo Fisher |
| ERCC RNA Spike-In Mix | Thermo Fisher |
| USB Dithiothreitol (DTT, 0.1 M) | Thermo Fisher |
| RNase inhibitor | Takara Bio |
| dNTP Mix (dATP, dCTP, dGTP, and dTTP, each at 10 mM) | Thermo Fisher |

| | |
|---|---|
| SpeedBeads magnetic carboxylate modified particles | Merck |
| Peg8000 | Sigma Aldrich |
| TAPS 0.2 M buffer soln., pH 8.5 | Thermo Fisher |
| 0.5 M EDTA, pH 8 | Sigma Aldrich |
| Sodium Chloride solution 5 M | Invitrogen |
| Ultra Pure TrisHCl 1 M pH 8 | Invitrogen |
| Qubit DNA HS | Thermo Fisher |
| Illumina compatible barcodes | IDT (see Supplementary Table 6 for barcode sequences) |

### Oligos:

| | | |
|---|---|---|
| Oligo-dT: | IDT | AAGCAGTGGTATCAACGCAGAGTACTTT<br>TTTTTTTTTTTTTTTTTTTTTTTTTTT<br>(N1:34333300)(N2:25252525) |
| IS_PCR | IDT | 5'-AAGCAGTGGTATCAACGCAGAGT-3' |
| TSO | IDT | 5'-AAGCAGTGGTATCAACGCA<br>GAGTACATrGrG+G-3' |
| ME-A | IDT | 5'-TCGTCGGCAGCGTCAGATGTG<br>TATAAGAGACAG-3' |
| ME-B | IDT | 5'-GTCTCGTGGGCTCGGAGATG<br>TGTATAAGAGACAG-3' |
| ME-Rev | IDT | 5'-/5Phos/CTGTCTCTTATACACATCT-3' |

ADA2_WT IDT /56-FAM/TGGAGGATT/ZEN/ATCGGAAGCGGGTG/3IABkFQ/

ADA2_Mut/WT_Fixed IDT /5HEX/TGGAGGACT/ZEN/ACAGAAAGCGGGTG/3IABkFQ/

ADA2 fwd: GGTGAGGAATGTCACCTACA

ADA2 rev: CATCAAACTCAGTGACGTTC

### 2. RNA preparation

Full length mRNA-sequencing is based on Smart-Seq2 protocol [32, 33]. Lysis plates containing cells were thawed and primer annealing was performed for 3 min at 72°C. 3 uL of reverse transcription mix (5x Reverse Transcriptase buffer (1 uL), Maxima H minus Reverse Transcriptase (0.05 uL), RNase Inhibitor (0.125uL), 100mM DTT (0.25 uL), 5M Betaine (1 uL), 1M $MgCl_2$ (0.03 uL), 100uM TSO (0.05 uL), $H_2O$

0.495) was added to each well, and reaction occurred at 42°C for 90 min, then heat inactivation at 70°C for 5 min. Next cDNA pre-amplification was performed by adding 7 uL of mastermix [2X Kapa HiFi HotStart ReadyMix (6 uL), 10 uM IS_PCR primer (0.12 uL), Lambda exonuclease (0.05625 uL) and $H_2O$ (0.8237 uL)]. PCR program was 37°C for 30 min, 95°C for 3 min, 22 cycles of 98°C for 20s, 67°C for 15s, 72°C for 4 min, then final elongation at 72°C for 5 min.

At this stage primers are removed by SPRI bead cleanup (prepared as here https://openwetware.org/wiki/SPRI_bead_mix#Ingredients_for_50_mL_2) at a ratio of 0.7:1. Concentration of independent wells is measured with Qubit DNA HS kit, and wells are diluted to 0.15 ng/uL.

**3. Library preparation**

Tagmentation was performed on the diluted cDNA, by adding 1 uL cDNA to 1.5 uL tagmentation mix (Tn5 (2.6mg/mL purified psfTn5-c006, Addgene plasmid #79107, loaded with standard Illumina Tn5 adapters (Meds A, MedsB, MedsRev))) (0.250 uL), 5X TAPS-PEG (Buffer is is 8% PEG, 5mM MgCl2, 10mM TAPS) 0.5 uL), H2O (0.750 uL)) and incubate for 10 min at 55°C. Then the reaction was stopped and transposome stripped of cDNA by adding 0.1% SDS (1 uL) and incubating for 10 min at 55°C. 7 uL barcoding mix was added (5x buffer (2.5 uL), 10mM dNTP (0.3 uL), 10% Tween (0.15 uL), Kapa HiFi (0.2 uL), H2O (3.85 uL) and 2 uL primer mix at 3.75 uM/primer. PCR program was as follows: 72°C for 3 min, 95°C for 30s, then 12 cycles of 95°C for 15s, 55°C for 30s, 72°C 45s, then final elongation at 72°C for 5 min.

Library was pooled and cleaned up 2x with 0.9:1 ratio of SPRI beads.

Libraries were sequenced on a Novaseq 6000. Add sequencing facility info here.

**4. qPCR**

For quantitative analysis of the two different alleles, 1uL of the diluted cDNA is amplified with Ada2 specific primers in the presence of WT or mutated and edited probes that are attached to different fluorophores. Reaction conditions were as follows: 2X Kapa HiFi HotStart ReadyMix (2.5 uL), 10uM forward and reverse primers (0.05 + 0.05 uL), 10 uM WT probe (0.05 uL), 10 uM Edited probe (0.05 uL), H2O (1.3 uL). PCR program was 95°C for 1 min, 35 cycles of 95°C for 15 s, 63°C for 45 s.

**5. Analysis of scRNA-seq data**

Cutadapt[34]was used to trim RNA sequence reads from adapters and low-quality bases. STAR[35]was used to align to hg38, with ERCC reads added. Picard[36]was used to remove duplicate reads. HTSeq[37]was used to summarize read counts. Cells with less than 20000 reads or 500 features were filtered out, as well as those with ACTB expression less than 0.01 quantile of the normal distribution. Seurat[38]was used to process the count data. Shortly, data was log normalized, 2000 variable features were found and the data were scaled; this was done separately per condition. Pathway analysis on scRNA-data was done as follows. FindMarkers function of Seurat was used to find markers between the two conditions of interest, with logfc.threshold=0. The resulting table was arranged based on avg_log2FC. The fgsea[39]package was used to read in the C7 and hallmark pathways from the Molecular Signature Database /Subramanian, Tamayo, et al. (2005, PNAS) and one or more of the following as appropriate: Liberzon, et al. (2011, Bioinformatics) and to perform gene set enrichment analysis based on the ordered avg_log2FC from FindMarkers and minSize=15 and maxSize=500.

**STAR-Fusion**[40] was used to predict fusion transcripts from the single cell data. Pseudobulk samples were prepared by merging fastq files per condition (unedited, edited +/- NHEJ inhibition, per individual), and run with default parameters. The deconvolved abridged fusion predictions were filtered by removing all fusions that were categorised as neighbouring by STAR-Fusion annotation.

To detect possible chromosomal loss due to editing all bam files from two individuals, a patient and a healthy donor were merged to create pseudobulk bam files using samtools[41] . cellsnp-lite[42] was used to genotype the samples, based on the 1000G phase 1 SNPs (https://www.internationalgenome.org/category/variants/), with settings minMAPQ=20, minLEN=30, UMItag=None, p=20, -I P, countORPHAN, exclFLAG=UNMAP, SECONDARY, QCFAIL, DUP. The output was filtered for GT="het" & INFO/AD[0]>1 using bcftools [43]. The vcf files were uploaded to Michigan Imputation Server[44], where the 1000G phase3 30x panel was chosen as a reference for imputation and Eagle2.4 was chosen for phasing. The imputed files were filtered with bcftools for R2>0.3 and TYPE="snp. Then cellsnp-lite was run on all individual cells against the imputed file for each sample, with the same parameters as before, except minCOUNT=10 and minMAF=0.2. The haplotype ratios were calculated and cells with only one haplotype on the q arm of chr22 were considered to have lost one chromosome.

#### 6. Analysis of qPCR data

qPCR analysis was performed on the RFU values per allele. A cutoff of an RFU value 200 was determined to decide which allele (wild type, mutated or edited) was being expressed in each cell.

### Mass spectrometry

#### 1. Sample preparation

T cells from three DADA2 patients and healthy donors were cultured as previously described and nucleofected on day 5 of the pipeline, where 1M cells/sample were mock nucleofected or ADA2-

edited. Mock edited cells were treated with DMSO and edited cells with 0.5µM KU0060648, 0.6µM IDT Alt-R enhancer V2 or DMSO for the first 24h after nucleofection. Cells were collected seven days after editing on day 12 of the pipeline, washed 2X with ice-cold PBS, pelleted and snap-frozen in liquid nitrogen. Pellets were stored at -80C until mass spectrometry sample preparation.

The samples were lysed in 8M Urea (#U5378-500G, Sigma Aldrich) in 100 mM ammonium bicarbonate (NH4HCO3 containing benzonase nuclease (415 units/ml, sc-202391, Santa Cruz Biotechnology). Total protein concentration was measured with Bio-Rad Protein Assay Dye (#5000006, Bio-Rad Laboratories). 50 µg of total protein was taken from each sample for reduction (5 mM dithiothreitol (#D9779, Sigma-Aldrich), alkylation (15 mM iodoacetamide (#122271000, Acros Organics), and overnight digestion with 2µg Trypsin/Lys-c Mix (V507A, Promega) at 37°C. After digestion, samples were acidified with 10% trifluoroacetic acid (TFA, #85049.051, VWR) and desalted with BioPureSPN PROTO 300 C18 Mini columns (#HUM S18V, Nest Group) according to manufacturer's instructions. After desalting the samples were dried in a centrifuge concentrator (Concentrator Plus, Eppendorf). The dried peptides were reconstituted in 40 µl buffer A (0.1% (vol/vol) TFA, 1% (vol/vol) acetonitrile (#83640.320, VWR) in HPLC grade water (#10505904, Fisher Scientific)).

For the DIA analysis the resuspended peptides were further diluted 1:60 in buffer A1 (1% formic acid in HPLC water). 20 µl was loaded into an Evotip (Evosep, Denmark) following manufacturer's instructions.

### 2. Mass spectrometry and analysis

The desalted samples were analyzed using the Evosep One liquid chromatography system coupled to a hybrid trapped ion mobility quadrupole TOF mass spectrometer (Bruker timsTOF Pro, Bruker Daltonics) (Meier, Brunner et al., 2018) via a CaptiveSpray nano-electrospray ion source (Bruker Daltonics). An 8 cm × 150 µm column with 1.5 µm C18 beads (EV1109, Evosep) was used for peptide separation with the 60 samples per day methods (21 min gradient time). Mobile phases A and B were 0.1 % formic acid in water and 0.1 % formic acid in acetonitrile, respectively. The MS analysis was performed in the positive-ion mode with dia-PASEF method[45] with sample optimized data independent analysis (dia) scan parameters. We performed DDA in PASEF mode from a pooled sample to be able to adjust dia-PASEF parameters optimally to these specific samples. To perform sample specific dia-PASEF parameter adjustment the default dia-short-gradient acquisition methods was adjusted based on the sample specific DDA-PASEF run with the software "tims Control" (Bruker Daltonics). The following parameters were modified for each sample type: m/z range; 429.2 – 1204.2, mass steps per cycle; 31 mean cycle time; 1.48 s. The ion mobility windows were set to best match the ion cloud density from the sample type specific DDA-runs.

To analyze diaPASEF data, the raw data (.d) were processed with DIA-NN v1.8.1 [46, 47] utilizing spectral library generated from the UniProt human proteome. During library generation following settings were used, fixed modifications: carbamidomethyl (C); variable modifications: acetyl (protein N-term), oxidation (M); enzyme:Trypsin/P; maximum missed cleavages:1; mass accuracy fixed to 1.5e-05 (MS2) and 1.5e-05 (MS1); Fragment m/z set to 100-1700; peptide length set to 7-30; precursor m/z set to 300-1600; Precursor changes set to 2-4; protein inference not performed. All other settings were left to default.

**3. Statistical analysis of the proteomics data**

The input file to further DIA data analysis was the DIA-NN Report.pg_matrix. For data pre-processing an in-house R-script was utilized. Raw intensity values were log2 transformed and median-normalized. Afterwards, missing values were imputed using QRILC imputation[48]. For sample group comparison, p-values were calculated with student's t-test using python package scipy[49], and adjusted using benhamini-hockberg method via statsmodels package[50]. Volcano plots were generated with bioinfokit[51] using q-value threshold of 0.01 and log2 intensity fold change thresholds of 1 and -1.

**Statistics:**

The following softwares were used for data analysis: QuantaSoft (Bio-Rad), FlowJo, Cutadapt 3.2, STAR 2.7.7a, HTseq 0.9.0, Picard 2.22.0, Seurat 5.0.1, FGSEA 1.20.1, STAR-Fusion V1.11.0., cellsnp-lite 1.2.3, bcftools 1.14. All of the statistics in the study were performed using GraphPad Prism 9.
